# Adaptation to volumetric compression drives hepatoblastoma cells to an apoptosis-resistant and invasive phenotype

**DOI:** 10.1101/2023.10.08.561453

**Authors:** Xiangyu Gong, Noriyoshi Ogino, M. Fatima Leite, Zehua Chen, Ryan Nguyen, Raymond Liu, Emma Kruglov, Kaitlin Flores, Aiden Cabral, Gabriel M. M. Mendes, Barbara E. Ehrlich, Michael Mak

## Abstract

Liver cancer involves tumor cells rapidly growing within a packed tissue environment. Patient tumor tissues reveal densely packed and deformed cells, especially at tumor boundaries, indicative of physical crowding and compression. It is not well understood how these physical signals modulate tumor evolution and therapeutic susceptibility. Here we investigate the impact of volumetric compression on liver cancer (HepG2) behavior. We find that conditioning cells under a highly compressed state leads to major transcriptional reprogramming, notably the loss of hepatic markers, the epithelial-to-mesenchymal transition (EMT)-like changes, and altered calcium signaling-related gene expression, over the course of several days. Biophysically, compressed cells exhibit increased Rac1-mediated cell spreading and cell-extracellular matrix interactions, cytoskeletal reorganization, increased YAP and β-catenin nuclear translocation, and dysfunction in cytoplasmic and mitochondrial calcium signaling. Furthermore, compressed cells are resistant to chemotherapeutics and desensitized to apoptosis signaling. Apoptosis sensitivity can be rescued by stimulated calcium signaling. Our study demonstrates that volumetric compression is a key microenvironmental factor that drives tumor evolution in multiple pathological directions and highlights potential countermeasures to re-sensitize therapy-resistant cells.

**Significance statement:** Compression can arise as cancer cells grow and navigate within the dense solid tumor microenvironment. It is unclear how compression mediates critical programs that drive tumor progression and therapeutic complications. Here, we take an integrative approach in investigating the impact of compression on liver cancer. We identify and characterize compressed subdomains within patient tumor tissues. Furthermore, using in vitro systems, we induce volumetric compression (primarily via osmotic pressure but also via mechanical force) on liver cancer cells and demonstrate significant molecular and biophysical changes in cell states, including in function, cytoskeletal signaling, proliferation, invasion, and chemoresistance. Importantly, our results show that compressed cells have impaired calcium signaling and acquire resistance to apoptosis, which can be countered via calcium mobilization.

## Introduction

Compressive stress is a crucial player in tumorigenesis and the progression of many cancers. Evidence of compressive stress caused by tumor growth has been found in several tumor types^1–4^. Additionally, in inflammatory tissue sites, biomolecular concentration is often elevated in the interstitial space, which increases the extracellular osmolarity^5,6^ and imposes osmotic pressure on local cells. Cancer cells also efflux sodium to balance intracellular tonicity under solid stress as a survival strategy^7^. Here, we focus on the impact of volumetric compression on liver cancer progression and phenotypic transitions. Liver cancer often arises as a consequence of chronic liver diseases, pathological conditions that are associated with altered biophysical properties of the tissue^8,9^. The implication of an altered mechanical environment, especially under compressive stress, in liver cancer progression is not well understood.

The tumor microenvironment (TME) is mechanically dysregulated with tumor growth, matrix stiffening, and interstitial fluid pressure in the TME leading to increased compressive stresses on cells. *In vitro* multicellular tumor spheroid (MCTS) models showed that growth in a confined space induces solid compressive stress in the constitutive cells^10–17^. A previous study demonstrated that planar mechanical compression enhanced breast cancer cell invasion by upregulating cell-substrate adhesion^18^. It was also shown that osmotic compression causes adipocytes to dedifferentiate into stem-like cells, which then differentiate into myofibroblasts and further facilitate breast cancer development^19^. This study provides evidence for indirect osmoregulation in cancer progression. On the other hand, *in vitro* studies combined with numerical models showed that mechanical stress induced by elevated osmotic pressure inhibits tumor growth^20^ but enhances the chemoresistance of cancer cells^21^. A recent study also showed that osmotic pressure could suppress cancer cell dissemination in a collagen matrix^22^. A study based on a mouse model showed that a persistent mechanical pressure applied to a healthy colon was sufficient to trigger tumorigenesis via β-catenin signaling^23^. These studies support that compressive stress can impact tumor and TME phenotypes in multiple ways. The cytoskeleton, a major component to facilitate cancer cell migration and mechanotransduction, is often associated with phenotypic changes in cancer^24^. For instance, altered mechanosensing in the cells can trigger tumorigenesis via YAP/TAZ mechanotransduction^25^. Small GTPases, especially RhoA and Rac1, are crucial cytoskeletal mediators during development and cancer progression. We recently identified subpopulations of liver cancer cells that use different cytoskeletal modes to invade into the adjacent normal^26^. How volumetric compression drives the cytoskeletal responses in the cells and the implication of mechanosensing and cell survival are unknown.

Regulation of critical liver functions, including cell proliferation, secretion, and apoptosis, depends on calcium^27^. Calcium channels inositol 1,4,5-trisphosphate (InsP3) receptors (ITPRs) are major players in intracellular calcium signaling in both normal and pathological hepatocytes^27–30^. The crosstalk between mitochondria and endoplasmic reticulum (ER) through ITPRs, leading to calcium flux, has been identified as a crucial factor in determining senescence^31^ and apoptosis^32^ - two important targets in cancer. Some mechanical cues, such as flow, stretching, and substrate stiffness, can regulate calcium signaling to dictate cell functions and cancer progression^33–36^. Intracellular calcium concentration has also been related to cell volume^37,38^. However, whether and how volumetric compression regulates cancer cell survival via calcium channels and intracellular calcium signaling is still poorly understood.

In this study, we report and characterize the existence of spatially distributed mechanically compressed subpopulations of liver cells in hepatocellular carcinoma (HCC) patients. We further experimentally investigate alterations and transitions in cell states (both molecular profiles and biophysical function) induced by conditioning liver cancer cells (HepG2 cell line) under volumetric compression over multiple days. We focus on the regulations of cytoskeletal and biophysical cell phenotypes, calcium signaling, and their interplay toward the emergence of two cancer hallmarks - cell invasion and death resistance^39^.

## Results

### Volumetric compression signals in patient liver tumors

In solid tumors, various forces are involved in cancer progression. In the liver, tissue stiffening due to liver fibrosis has been shown to regulate cell behavior mechanically and contributes to HCC progression^8,9^. Here we evaluate whether another mechanical factor - compressive stress - can be generated in liver tumors that may alter cell structure and functions. Evaluation of histological samples from multiple HCC patients revealed that a subpopulation of liver cells or HCC cells are subject to compression (Fig. 1a). These compressed cells are often seen around fibrotic regions and exhibit distinctly smaller and elongated morphology (Panels 1 and 5), compared to nearby non-fibrotic counterparts within the same tumor (Panels 2 and 6). It appears that the expansion of the cell-packed tumors built up the pressure so that the constitutive cells near the fibrotic regions were pressed against regions with stiff, dense collagen features. Fat droplets in steatotic regions of tissue of HCC patients (SH-HCC) also lead to the deformation of the cells and cell nucleus (Panel 7), a morphological change that was shown to regulate cell mechanotransduction^40^. Furthermore, HCC tumors can grow into the adjacent normal hepatocyte regions (peritumor regions), which would exert significantly high compression on the normal cells. This invasive effect is observed when comparing Panel 4 (tumor-free normal liver region) with Panel 3. At the interface of the tumor and the peritumor normal tissue, the hepatocytes are dramatically deformed, featured with narrowed sinusoidal space, packed nucleus, and smaller and elongated cell body. In-house image analysis codes were built to further characterize the density and shape of cell nuclei in larger regions of the HCC histology samples. In cases where the tumor grows against normal cells, we found that nuclear density, which is indicative of cell packing, was highest near the edge of the tumor boundary (Figure 1b, Patient 1). This packing decreases further away from either side of the tumor boundary. In addition, nuclear area is the smallest near the tumor boundary and gradually increases moving away from the tumor boundary. Increased nuclear packing and decreased area are also present at the boundary of tumors growing against a fibrotic capsule (Figure 1b, Patient 2). Together, these metrics demonstrate the presence of compressive forces against the tumor boundary.

**Fig. 1.**
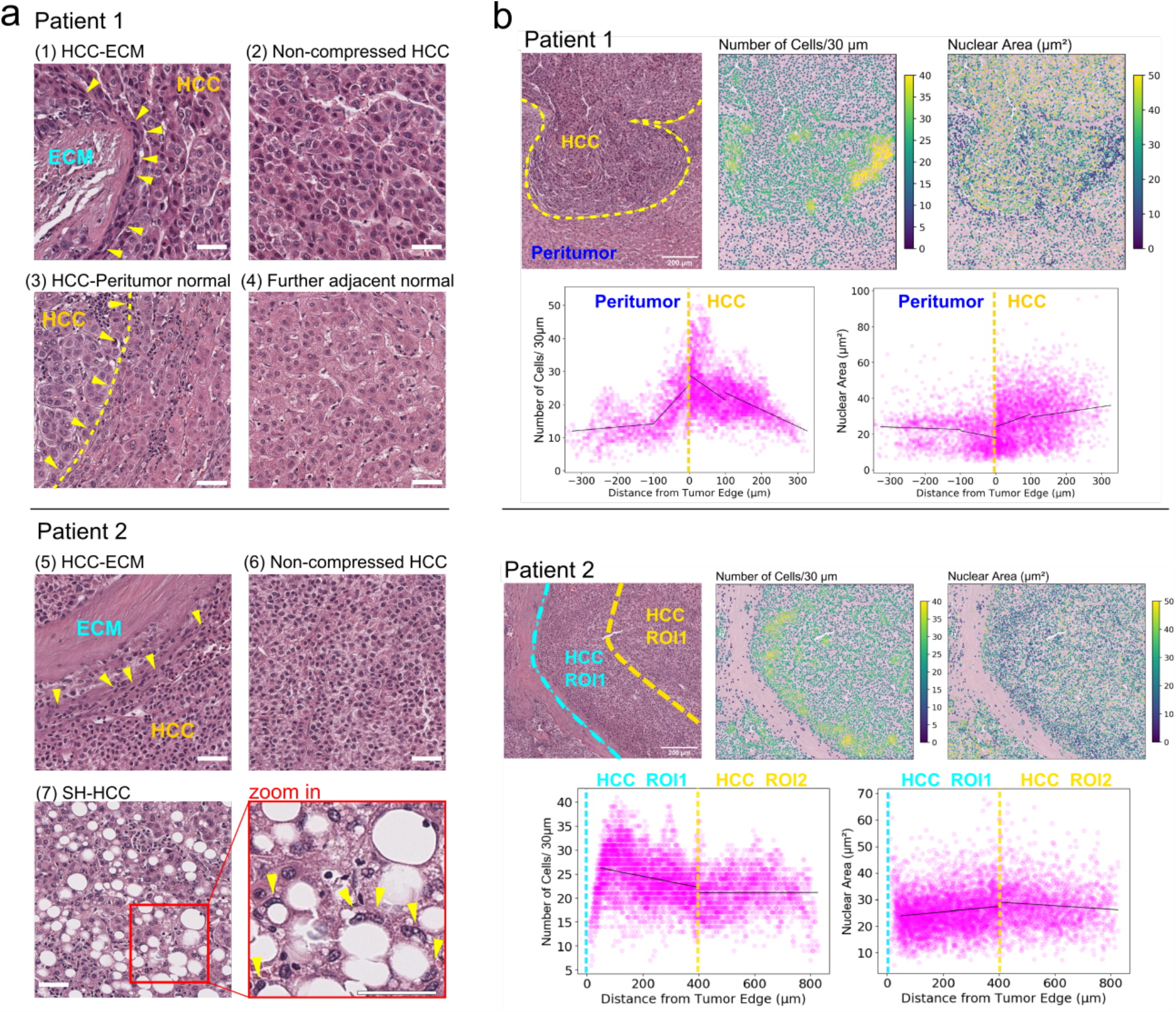
Mapping the compression signatures in liver cancer. (a) H&E staining of HCC samples shows locally compressed regions of cancer cells or normal adjacent liver cells due to hyperproliferation, fibrosis, and steatosis (SH). In the compressed regions, the cells exhibit highly deformed cell bodies and nuclei. Scale bars in (1-4): 50 μm, and in (5-7): 100 μm. Yellow arrowheads indicate the deformed cell nuclei. The dotted line in (6) indicates the boundary between the HCC tumor and the peritumor normal liver cells. (b) Quantification of compression in HCC samples. Tumor regions growing against either normal adjacent tissue or fibrotic capsule (dense fibrotic region surrounding tumor) demonstrate high nuclear density and lower nuclear area near the tumor boundary. Scatter plots indicate nuclear density and the area of cells within the tumor boundary with respect to the tumor edge. The dotted yellow lines represent tumor boundaries. In Patient 2, the region (ROI1) between cyan and yellow lines has strong gradients of nuclei density and nuclei area. Black lines in the scatter plots indicate piecewise linear regression. Scale bars: 200 μm. Images are representative of two HCC patients evaluated.

### Hyperosmolarity-induced volumetric compression drives morphological and phenotypic changes in hepatoblast-like liver cancer cells

The morphological analysis showed that in patient-derived tissues both HCC cells and hepatocytes are subjected to compressive stress. To observe the outcome of a persistent compression and restriction of cell volume in isolated liver cancer cells, we introduced volumetric compression for at least 5 days by hypertonic culture conditions at different levels of osmolarity (Fig. 2a). We chose the liver cancer cell line HepG2 as the model cell type given that the cell line recapitulates characteristics of hepatoblasts as well as epithelial-like HCC^41^. Several previous studies showed that acute osmotic stress induces cell death^42,43^. Here we found HepG2 cells at all different compression levels after long-term treatment (5 days) not only survived better than under control conditions but also underwent a drastic morphological change (Fig. 2b). Although the hepatoblast-like cancer cells (HepG2 cells) tended to grow into multicellular colonies on 2D surfaces with no indication of outward cell migration, the compressed counterparts grew into flat colonies with reduced height (Fig. 2c,d). Under high compression (5 days in 4% PEG), the cells grew into a flattened monolayer and exhibited a dramatic morphological change. Compared to cells maintained under isotonic conditions, the compressed cells had a larger spreading area and turned from the typical round, cobblestone-like cells into highly elongated spindle cells. Also, over the course of 5 days, unlike the isotonic cells, the compressed cells did not exhibit severe cell death (Fig. 2e) or activate apoptosis labeled by cleaved caspase 3/7 (Fig. 2f,g). An apoptosis-inducing drug staurosporine (STA), a potent protein kinases inhibitor, which significantly triggered cell apoptosis in the isotonic condition, failed to trigger apoptosis in the compressed cells.

**Fig. 2.**
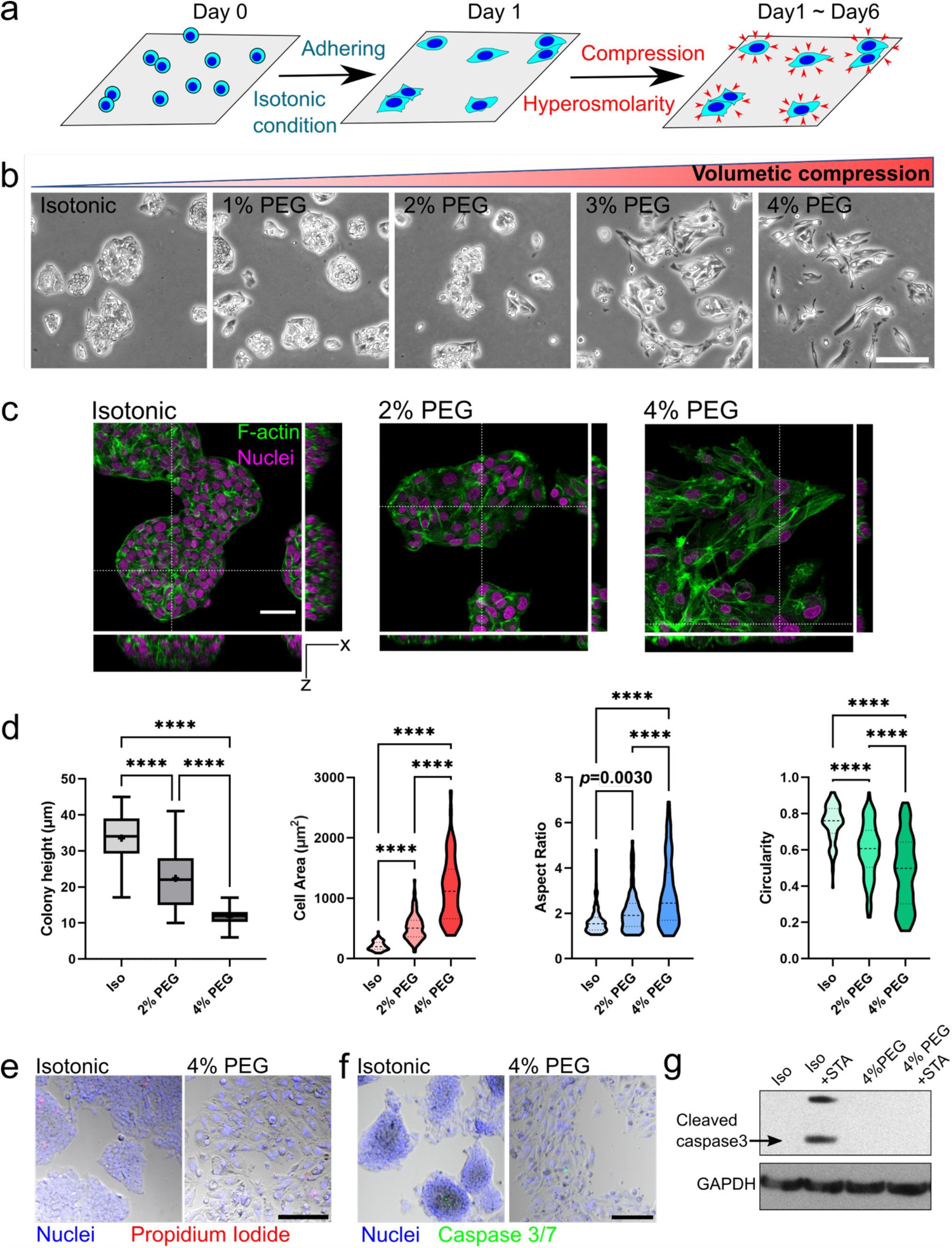
Volumetric compression induced a drastic morphological change in HepG2 without triggering apoptosis. (a) Schematic of compression method. (b) HepG2 grew into less circular and more irregular multicellular colonies over five days under compression. (c) Fluorescence staining of the day-5 colonies under isotonic, 2% PEG and 4% PEG conditions revealed: (d) a reduction in colony height under compression and a significant increase in cell spreading area and elongation. For colony height: n=36-44 colonies from N= 3 replicates. For cell area, aspect ratio, and circularity, n= 71-119 cells from N=3 replicates. Scale bar, 50 μm. (e) Cell viability of isotonic vs. 4% PEG compression. Scale bar, 100 μm. Cell apoptosis was characterized by cleaved caspase-3/7 labelled by (f) CellEvent, and validated by (g) immunoblot for cleaved caspase 3. Scale bar in (f), 200 μm. For multiple comparisons in (d) were made using one-way ANOVA with Tukey post hoc (\*\*\**p*<0.001, \*\*\*\**p*<0.0001).

Using bulk RNA-seq, we interrogated which signaling pathways were regulated and linked to the morphological changes of the long-surviving compressed HepG2 cells (Fig. 3). Compared to the non-compressed colonies grown, the compressed cells (5 days in 4% PEG) had a shifted transcriptional landscape (Fig. 3a). Consistent with previous studies^41^, expressions of liver-specific genes were enriched in hepatoblastoma HepG2 (Fig. 3b). Under compression, the cells lost their liver identities but acquired an epithelial-mesenchymal transition (EMT) phenotype (Fig. 3c). The primary upregulated pathways in the compressed cells included cytoskeleton rearrangement, adhesion, cell-matrix interactions, EMT, and anti-apoptosis (Fig. 3d). Notably, consistent with studies on other tissue types^19,23,44^, volumetric compression upregulated Wnt signaling in HepG2 cells (Fig. 3d) and triggered β-catenin translocation from cytoplasm membrane to the nucleus (Fig. S1). Major downregulated pathways were cell cycle-related, possibly due to the DNA damages (Fig. 3e). Altogether, the compressed cells appeared to adopt an invasive, but non-proliferative, phenotype.

**Fig. 3.**
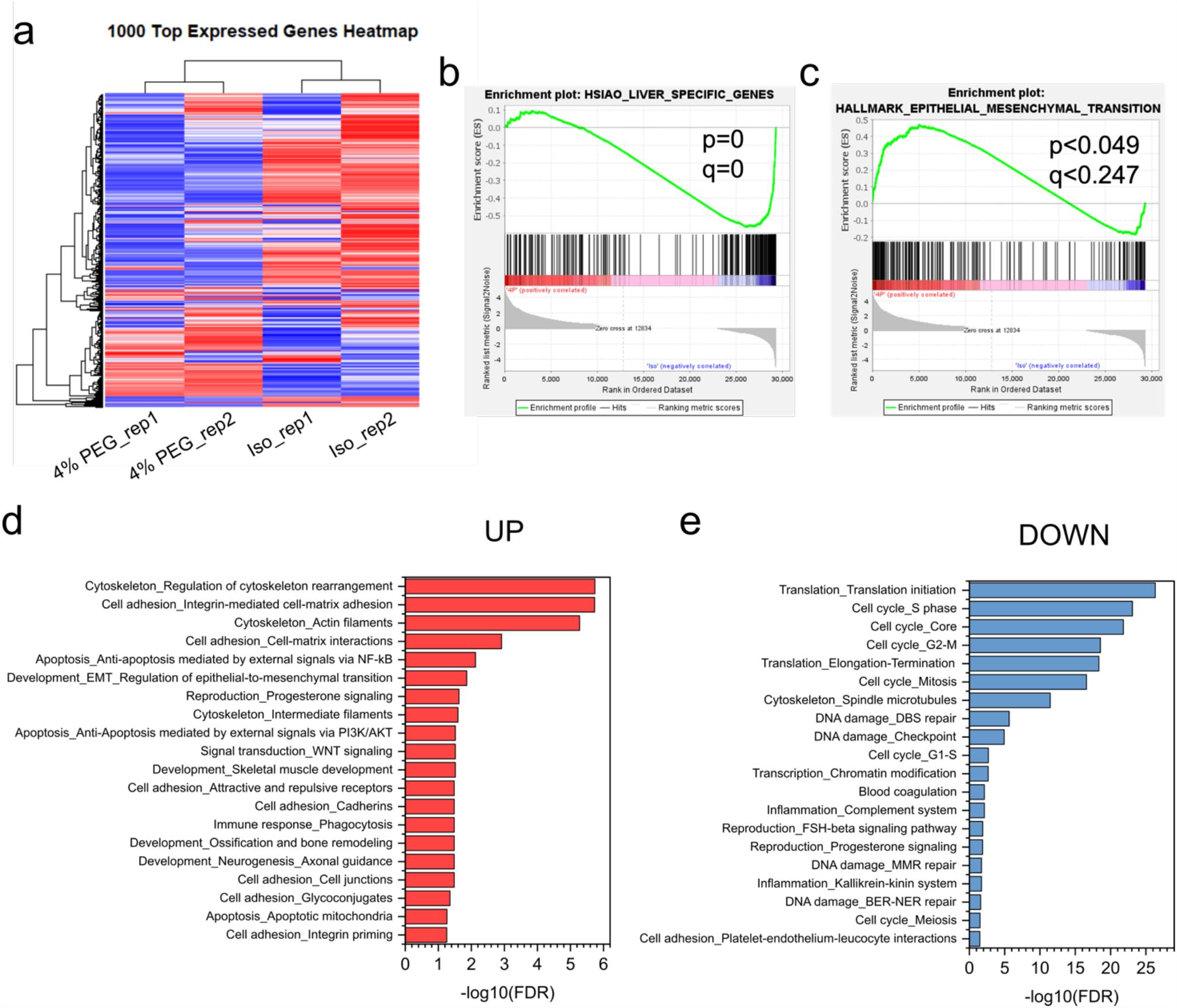
RNA-seq revealed a phenotype change in volumetrically compressed HepG2 cells. (a) The top 1000 regulated genes indicated a cell phenotypic transformation under compression. (b, c) Geneset enrichment analysis showed that hepatoblast-like HepG2 cells lost hepatocyte signatures and potentially underwent epithelial-to-mesenchymal transition (EMT). (d) Gene ontology analysis demonstrated the top 20 upregulated and downregulated cell behavior in compressed cells.

### Cytoskeletal dynamics mediate compression-induced cell spreading and survival phenotypes

Cytoskeleton regulation was involved in this dramatic morphological change in the compressed cells. Daily tracking of the compressed cells revealed a temporal characteristic of the cell morphology (Fig. 4a). Under compression, the cells started exhibiting a significant change in both spreading area and cell shape between Day 2 and Day 3. From Day 3, the cells became increasingly larger and spindly over time (Fig. 4b-d). In wound healing assays (Fig. 4e), compression rendered a comparable wound closure rate of the cells over a week, despite a significant halt in the cell proliferation measured by EdU-positive cells between Day 5 and Day 6 (Fig. 4f). This lack of proliferation implied that enhanced cell spreading under compression was the main driver for collective cell movement. With higher cell-substrate engagement and cell spreading, the compressed cells also exhibited higher YAP nuclear translocation (Fig. 4g), measured by the intensity ratio of nuclear YAP and cytoplasmic YAP in individual cells (Fig. 4h), as well as the percentage of the whole cell population with high YAP translocation (ratio>1) (Fig. 4i). Nuclear translocation of YAP is mechanosensitive and controlled by cell tension and nucleus stretching^45^. Under isotonic conditions, the cells in the colony core often had a low nuclear YAP signal compared to the cells at the colony edge, suggesting that the internal confinement during the tumor expansion reduced cell tension and thus downregulated the YAP activity. In contrast, under compressed conditions with high cell-substrate engagement, a larger subpopulation of cells maintained a high cell tension and higher YAP nuclear translocation. When compressing the cells on elastic polyacrylamide (PA) substrates with a physiological or pathological stiffness^46^, we still observed drastic phenotype changes, including enhanced cell spreading and YAP nuclear translocations (Fig. S3 a,b). This suggested that the compression-induced cell phenotype was substrate stiffness independent.

**Fig. 4.**
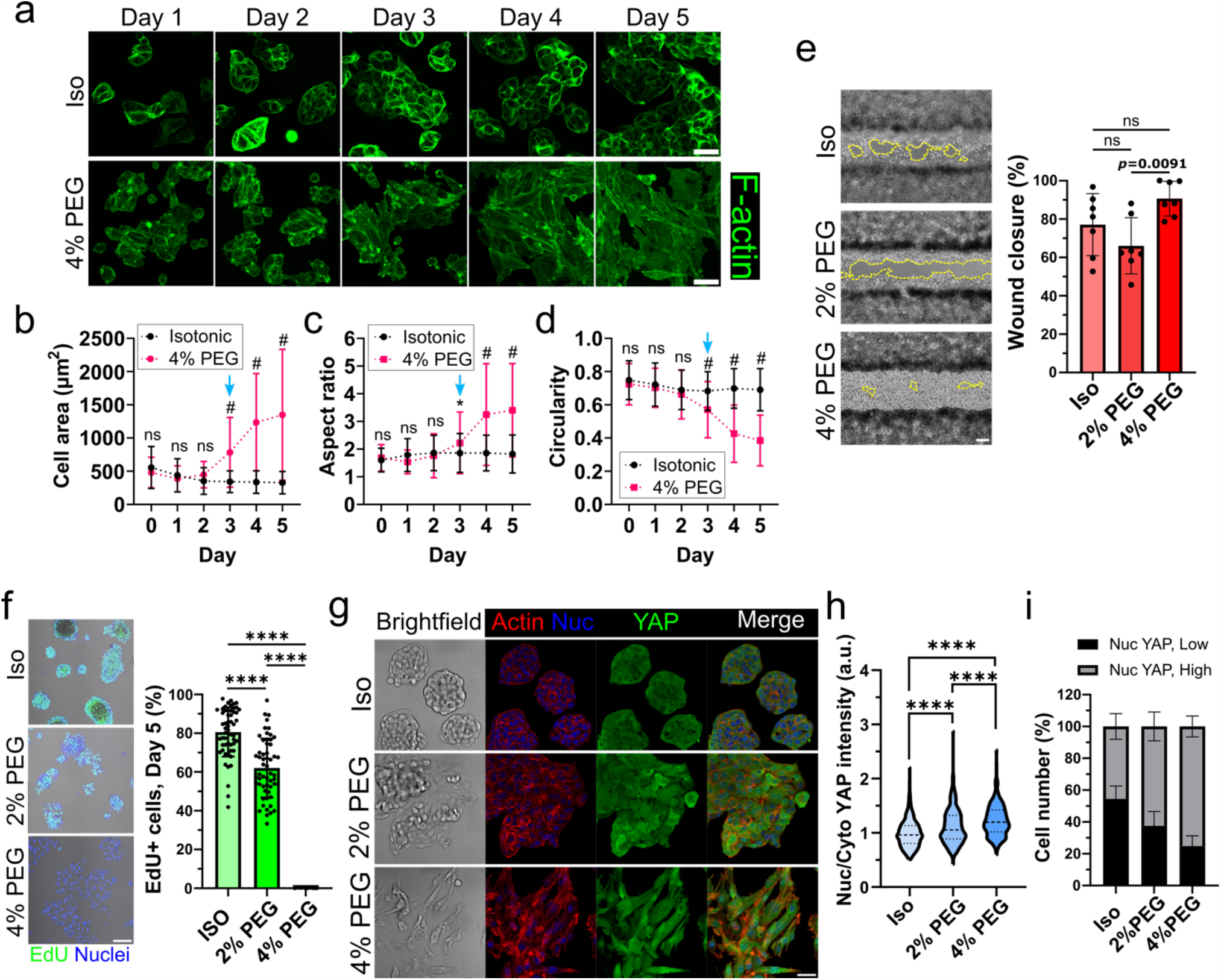
Temporal mechano-adaptation of HepG2 enabled cell-substrate engagement and wound healing despite a halt in cell proliferation. (a) Fluorescent imaging (F-actin) showing the cell morphology over the course of five days in the isotonic conditions vs. 4% PEG medium. Scale bars, 50μm. The cell morphology was depicted as (b) spreading area, (c) aspect ratio, and (d) circularity. n=119-369 cells for “Isotonic”, and n=91-148 cells for “4% PEG” on each day. (e) Wound healing assay measurements on Day 8. n=7 wounds from n=3 biological replicates each condition. (f) EdU staining comparing cell proliferation over 12 hours in the isotonically cultured cells and the 5-day compressed cells. The percentages were calculated based on n=55, 56, 8 colonies for Iso, 2% PEG, and 4% PEG conditions respectively, from N=2 independent experiments. Scale bars in (e) and (f), 200μm. (g) Immunofluorescent staining showing cell shape, actin organization, and YAP subcellular location. Scale bar: 50μm. The extent of YAP nuclear translocation was determined by (h) the fluorescent intensity ratio of YAP in nuclei and cytoplasm, and (i) the percentage of cells with high nuclear YAP vs. low nuclear YAP within instinct colonies. For (h, i) n=241-350 cells from N=3-4 individual colonies. The threshold of “high nuclear YAP” is “Nuc/Cyto YAP intensity > 1”. In (b-d), two-way ANOVA with Tukey post hoc was used to make comparisons between days and culture conditions. Here, only the statistical significances between culture conditions on each day were shown in the plot (*p=0.0202, #p<0.0001). Multiple comparisons in (e), (f), (h) were made using one-way ANOVA with Tukey post hoc (\*\*\*\**p*<0.0001).

To understand the cell spreading mechanism at the single cell level, we conducted time-lapse imaging on LifeAct-labeled HepG2 cells daily, after the regular culture condition was switched to the compression condition (4% PEG). Upon compression (Day 0 to Day 1, Supplementary Video 1), no significant morphological changes occurred, as supported by Fig. 4 a,b. Between Day 3 and Day 4, we started to observe the difference between isotonically cultured cells and the compressed cells, in terms of cell morphology and actin dynamic (Fig. 5a,b, Supplementary Video 2). In the isotonic condition, multicellular colonies, as a whole, were barely mobile, and there were no protruding cells growing out of the colonies (Fig. 5a). In contrast, the cells in a 4% PEG compressed colony gradually shifted their cortical actin into sheet-like extensions with edge ruffling (Fig. 5b) resembling lamellipodia. Small Rho GTPases RhoA and Rac1 are master regulators of the cytoskeleton^47^. Using G-LISA, we measured the activities of these two regulatory components. We found an increased activity of Rac1 and a decreased activity of RhoA in the highly compressed cells (Fig. 5c). The shifted ratiometric balance in RhoA and Rac1 was likely responsible for the newly formed lamellipodia. The regulation of this ratio shift was tested by inhibiting Rac1 or actin polymerization during the 5-day compression. We found that Rac1 inhibitors, NSC23766 (NSC) and EHT 1864 (EHT), or actin nucleator ARP2/3 inhibitor CK666 can suppress compression-enhanced spreading and elongation measured by cell area, aspect ratio, and circularity (Fig. 5d-g). Inhibiting cell contractility rendered a slimmer cell body thus smaller cell area, but it did not inhibit cell extension, suggesting the compression-driven protrusion was actomyosin independent. We found that stabilizing microtubules with paclitaxel rather than depolymerization with nocodazole (Noco) blocks the compression-driven cell spreading. This observation showed that microtubule dynamics also played a role in the compression-induced cell morphology shift.

**Fig. 5.**
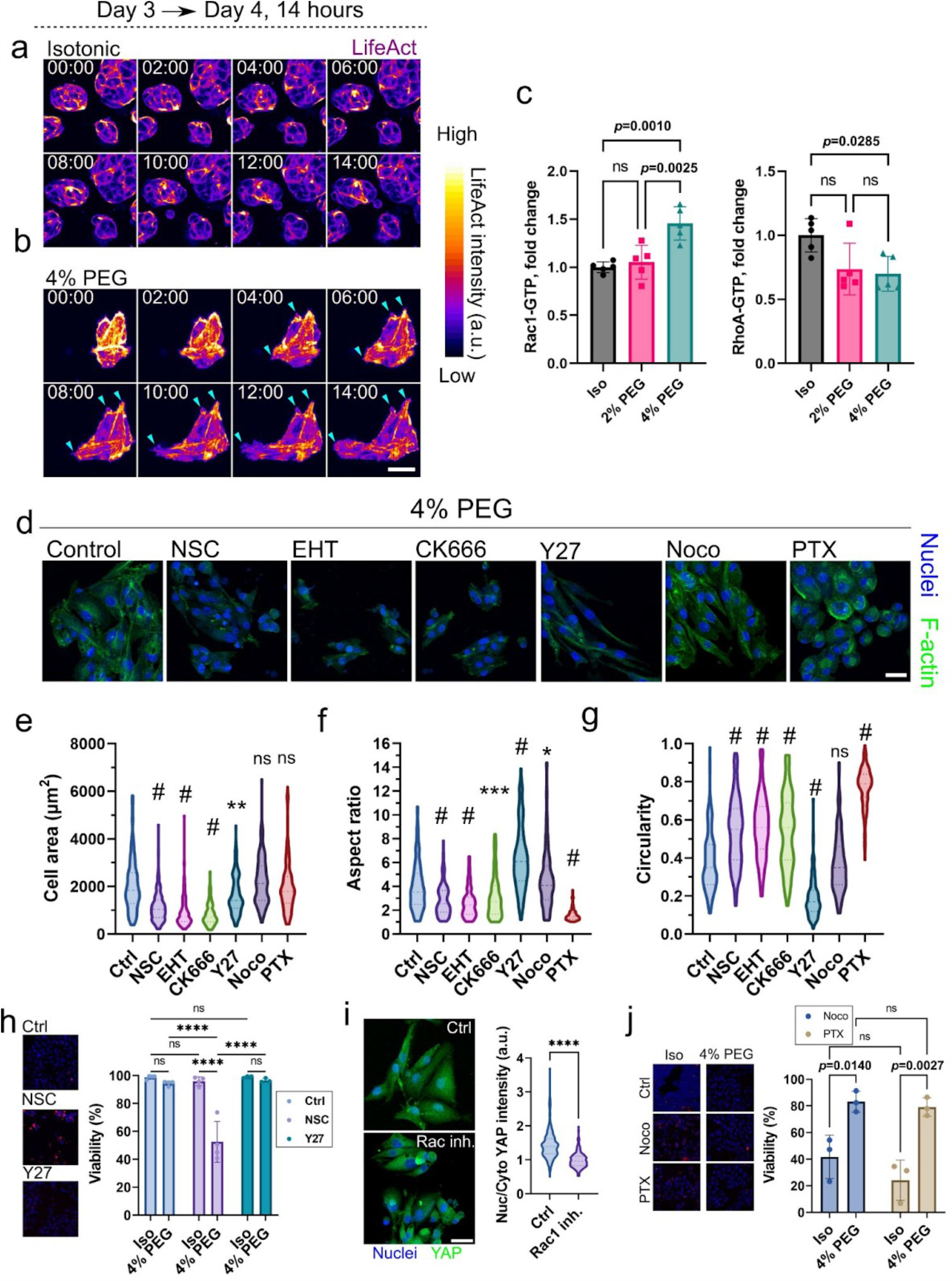
Cytoskeletal regulations are involved in cell spreading and survival under volumetric compression. Time-lapse imaging on LifeAct (14 hours) between Day 3 and Day 4 of the culture, showing cytoskeleton dynamics and cell morphology of HepG2 colonies expanding in the (a) isotonic condition and (b) 4% PEG condition. Scale bar, 50 μm. (c) GLISA assay was used to determine the activities of small Rho GTPases RhoA and Rac1. n=5 duplicate wells from N=2 biological samples. (d) F-actin staining showing cell morphology of 4% PEG compressed cells with the treatments of different inhibitors at Day 5 (Rac1 inhibitor: NSC and EHT; ARP2/3 inhibitor: CK666; ROCK inhibitor: Y27; Microtubule destabilizer: Noco; Microtubule stabilizer: PTX). Scale bar: 50 μm. The cell morphology was measured by (e) spreading area, (f) aspect ratio, and (g) circularity. For each condition, n = 109-223 cells from N = 3 biological replicates. Statistical comparisons were made between individual drug treatments and the control (CTRL), using ANOVA with Tukey post hoc. (\*\**p*=0.0037 in (e), \**p*=0.0263 in (f), \*\*\**p*<0.001, #*p*=0.0001). (h) Cell viability of isotonically cultured vs. 4% PEG compressed cells with Rac1 inhibition and ROCK inhibition at Day 5. N=3-4 biological replicates each. (i) Quantification of YAP nuclear translocation in the compressed cells after 5 days of Rac1 inhibition (Rac1 inh.), compared with compressed cells without drug treatments (Ctrl). n= 268 for control and 176 for Rac inhibition from N=3 biological replicates. Scale bar, 50 μm. (j) Comparing cell viability after 5-day treatment of the microtubule drugs (Noco and PTX) under isotonic vs. 4% PEG conditions. N=3-4 biological replicates each. One-way ANOVA with Tukey post hoc was used in (c) and (i) (\*\*\*\**p*<0.0001). For (h) and (j), two-way ANOVA with Tukey post hoc was used for the statistical analysis (\*\*\*\**p*<0.0001).

The viability of the cells was also monitored with drug screening (Fig. 5h), and we found Rac1 activity, but not cell contractility, is crucial for cell survival under compressive stress. We further found the inhibition of Rac1 in the compressed cells also reduced the nuclear translocation of YAP (Fig. 5i). Disrupting microtubules impairs mitosis to induce cell death^48^. Remarkably, despite the different effects on cell morphology, with 5-day treatments of both microtubule disruptors, we found much higher cell viability under compression compared to the cells growing in the isotonic condition (Fig. 5j). Altogether, we demonstrate that the cells rearrange the cytoskeleton to survive when adapting to severe volumetric compression.

### Volumetric compression regulates the expression of InsP3 receptors (ITPR) and alters intracellular calcium signaling

Calcium signaling is a vital physiological feature that impacts functions, including cell contraction^49^, proliferation^50^, and apoptosis^32^. We next interrogated whether intracellular calcium signaling and related calcium channels in the cells are altered when cells adapt to compression. The RNA-seq data revealed that volumetric compression-induced cell phenotype was associated with dysregulation of ITPRs (Fig. 6a). At the transcriptional level, the expression of ITPRs was altered as a function of time in culture (FigS. 3a), indicating a gradual shift of the cell phenotype as a response to the conditions of compression. By analyzing the data from The Cancer Genome Atlas (TCGA), we noticed a link between the regulations of ITPRs and liver cancer prognosis (FigS. 3b), showing the upregulation of ITPR3 was associated with poor prognosis^30^. To further verify the regulations of the ITPRs in the compressed cells, we quantified the expression of all three types of ITPRs at the protein level (Fig. 6b). Overall, the compressed cells showed slight upregulation of ITPR1, downregulation of ITPR2, and significantly increased expression of ITPR3 (Fig. 6c). This dysregulation directly impacted the intracellular calcium response upon ATP stimulation (Fig. 6d-f). Despite the overexpression of ITPR3 and the enhanced proximity of ITPR3 and mitochondria (Fig. 6 h,i), upon the stimulation of compressed cells with extracellular ATP, a standard agonist to invoke intracellular calcium release, the cells also exhibited reduced calcium uptake into mitochondria (Fig. 6g). To test whether the lower amplitude of intracellular calcium signaling induced by a calcium mobilizing agonist (ATP) was due to reduced calcium concentration inside the ER, calcium release to empty the ER was done using thapsigargin (TG) in both isotonic and compressed conditions. Upon the TG treatment (1μM), a high calcium signal was detected in the cells under all conditions (Fig. 7a) showing that the ER calcium load was similar and sufficient to produce a surge in calcium when stimulated in both isotonic and compressed cells. The compressed cells released even more calcium from the ER, which suggests the long-term compression did not affect intracellular calcium storage but instead reduced the calcium release via intracellular calcium channels. The BCL-2 family proteins are proteins that bind to ITPRs and suppress calcium release^51^. Our RNA-seq data indeed show an upregulation of anti-apoptosis BCL-2 family members (Fig. 6j), MCL-1 and Bcl-X (BCL2L1) suggesting an additional way for suppression of calcium release from the ER.

**Fig. 6.**
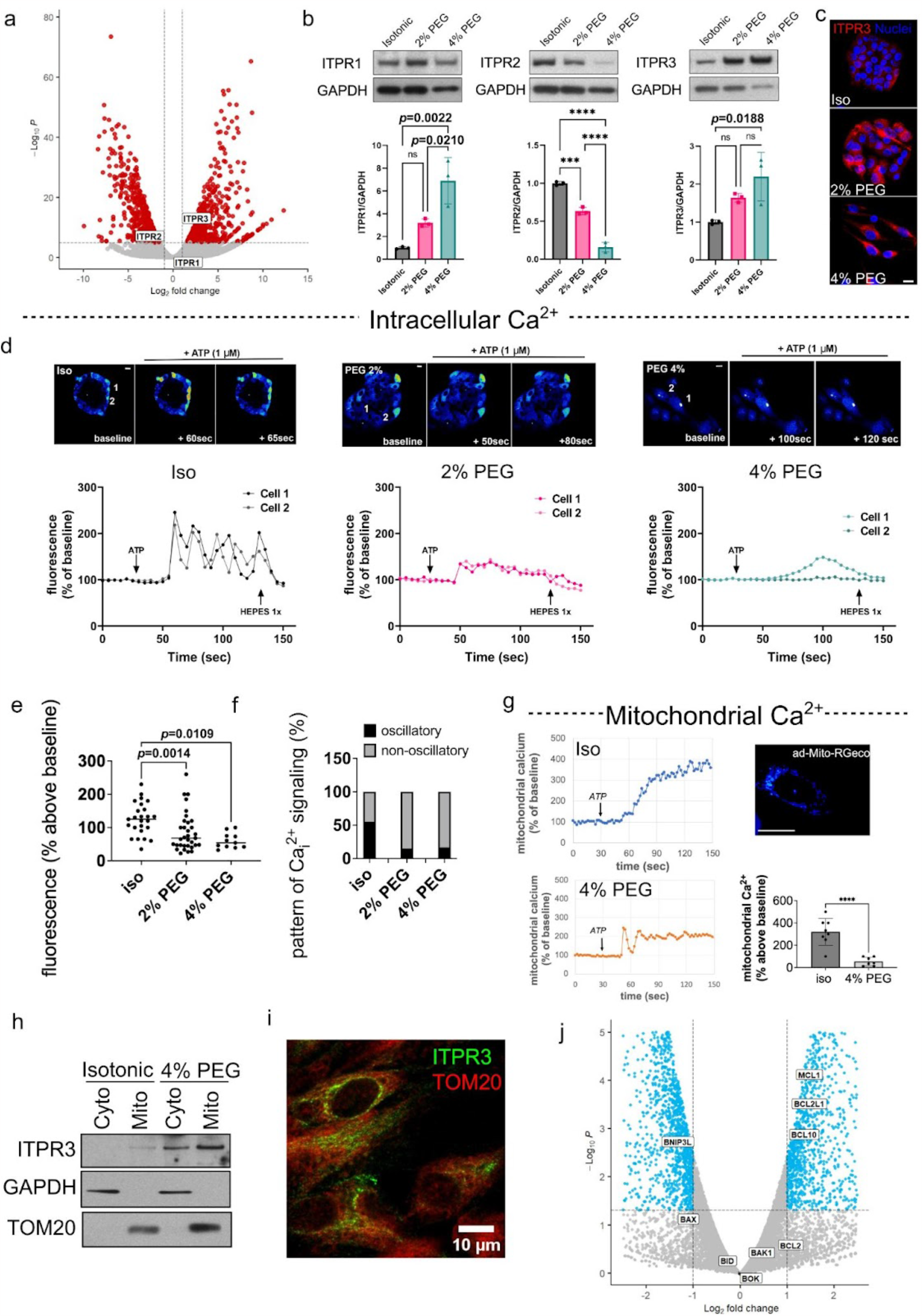
Volumetric compression regulates expressions of InsP3 receptors (ITPRs) and alters calcium signaling. (a) Volcano plot showing the regulations of ITPRs from the RNA-seq analysis. (b) Immunoblots and (c) fluorescent staining confirmed the expression level of ITPRs in the compressed cells compared to the cells cultured in the isotonic condition. (d) Intracellular calcium signaling upon the ATP (1μM) stimulation showed less calcium response in the compressed cells, in terms of both (e) amplitude and (f) frequency of oscillation. (n=25, 33, 11 from N=3 replicates each condition) (g) Mitochondrial calcium signaling showed that the compressed cells had less calcium response upon ATP stimulation. (h) Mitochondria extraction assay showed more abundant ITPR3, suggesting enhanced proximity between the mitochondria and ITPR3 in the compressed cells. This was further demonstrated in the immunofluorescent staining in (i). (j) RNAseq analysis summarizing the regulations of BCL-2 family members in the compressed cells. One-way ANOVA with Tukey post hoc was used in (b). One-way ANOVA with Bonferoni post hoc was used in (e). A two-tailed student’s t-test was used in (g) (\*\*\*\**p*<0.0001).

**Fig. 7.**
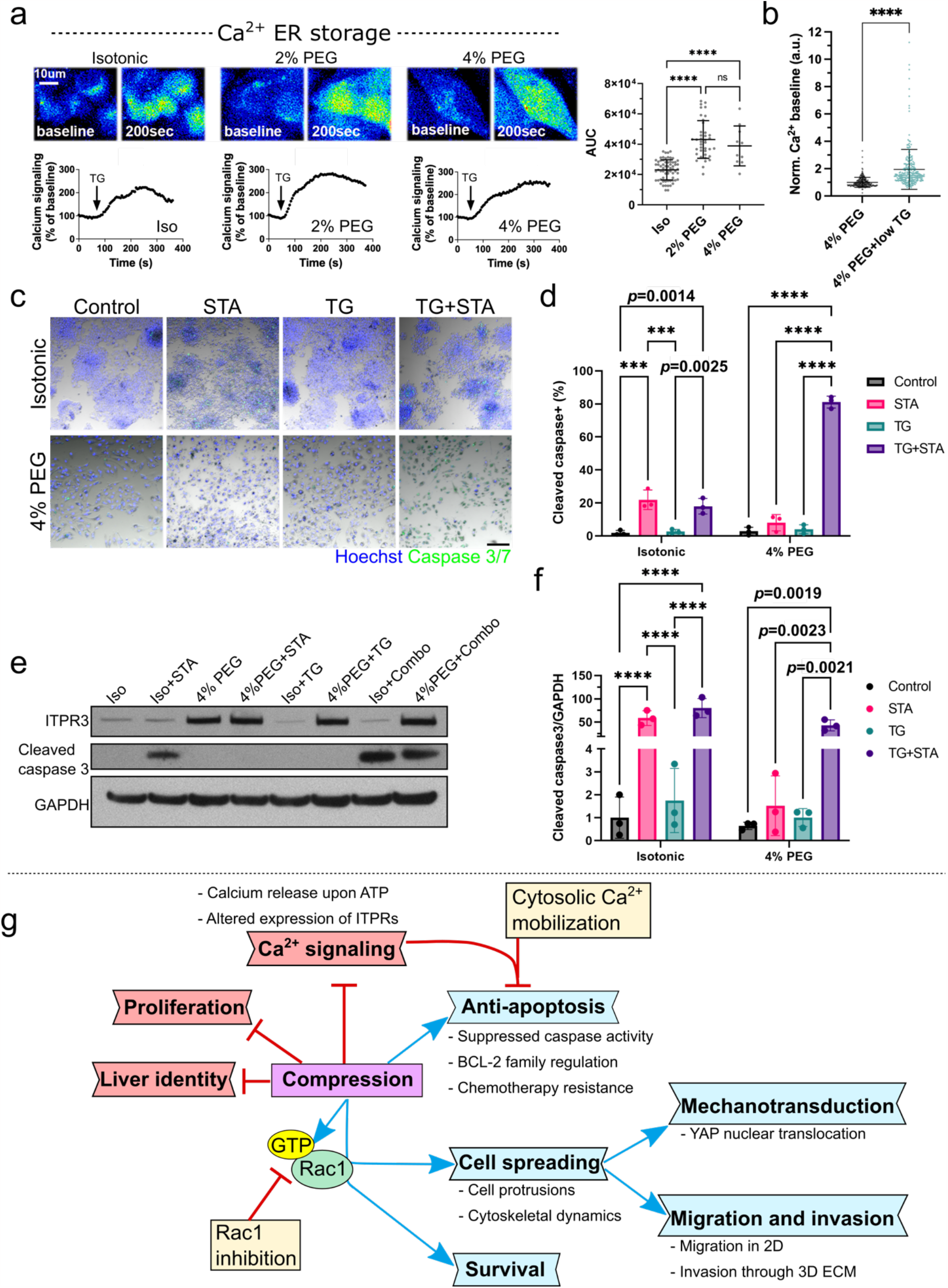
ER calcium leakage rescues compression-induced apoptosis resistance. (a) Intracellular calcium signaling (Fluo-4/AM) upon thapsigargin (TG) treatment (1μM). The corresponding measurements of calcium leakage from ER are represented as the area under the curve (AUC) (n = 66, 39, 10 cells from N = 2 coverslips for each condition). (b) Baseline calcium in the 4% PEG compressed cells with and without the overnight treatment of low-dosing TG (50nM), normalized by the mean of the 4% PEG control of each replicate. (n=287 cells for 4% PEG control, and 297 cells for the TG treated group, from N=3 biological replicates). (c) Cell apoptosis is indicated by Caspase 3/7 apoptosis assay under isotonic or 4% PEG conditions with STA and/or low-dosing TG treatment (50nM). The percentages of caspase 3/7 positive cells under different conditions are shown in (d) (N = 3 biological replicates). (e) Western blot further validates the protein level of ITPR3 and cleaved caspase 3 under the drug treatments. The fold changes of normalized production of cleaved caspase-3 under the treatments are shown in (f) (N = 3 biological replicates). In (a), one-way ANOVA with Tukey post hoc was used (\*\*\*\**p*<0.0001). In (b), two-tailed student’s t-test was used (\*\*\*\**p*<0.0001). In (d) and (f), two-way ANOVA with Tukey post hoc was used for the statistical analysis (\*\*\**p*<0.001, \*\*\*\**p*<0.0001). (g) Model of the adaptation of HepG2 in volumetric compression. Red boxes indicate suppressed phenotypes and blues boxes indicate the promoted phenotypes. Yellow boxes highlight the druggable interventions targeting the promoted phenotypes. Although hepatoblast-like liver cancer cells HepG2 retain strong liver identity, they exhibit rapid, unchecked colony expansion. Under long-term (at least 5 days) volumetric compression, the cells have significantly reduced proliferation, yet exhibit long survival without triggering apoptosis and become less sensitive to chemotherapy drugs. Over the course of surviving, the cells lose the liver phenotype, which is marked by the generation of Rac1-directed cytoskeletal shift and stronger cell-substrate engaging, which can be suppressed by Rac1 inhibition. This enhanced cell spreading leads to a monolayer colony growth and sustained high YAP nuclear translocation. The long-term compressed, non-apoptotic cells also have suppressed calcium signaling with an altered release from ER storage, which can be targeted by inducing calcium mobilization to reverse the anti-apoptosis phenotype of the compressed cells.

### Rescuing intracellular calcium signaling reverses apoptosis resistance in the compressed cells

In the previous section, we showed that the expression of BCL-2 family proteins is increased (Fig. 6j). These proteins are also known to regulate cell apoptosis via mitochondrial calcium^52,53^. We have shown that the compressed cells remained viable and did not respond to apoptosis even upon apoptosis induction by STA treatment (Fig. 2e-g). We next tested whether the attenuated intracellular calcium signaling in the compressed cells is responsible for the resistance to apoptosis. As ER calcium storage was unaffected (Fig. 7a), we reasoned that using a pathway other than ITPRs to mobilize the calcium from ER would restore the cells’ sensitivity to apoptosis.

We could mobilize calcium in the compressed cells by an overnight treatment of TG at a very low concentration (50nM), which was validated by the baseline calcium intensity (Fig. 7b). Cleaved caspase 3 was evaluated as the apoptosis marker in both imaging-based apoptosis assay (Fig 7. c,d) and immunoblotting (Fig. 7e,f). We showed that low-dosing TG treatment alone was not sufficient to trigger significant cell apoptosis in either the non-compressed or the compressed conditions. When treating the cells with the combination of TG and STA overnight, we indeed observed significant apoptosis in the compressed cells that did not respond to treatment by either of the drugs alone. Together, this result showed that low intracellular calcium in the compressed cells due to dysregulated ER-mitochondria communication was responsible for the non-apoptotic cell phenotypes. Moreover, a slight elevation of the resting calcium through leakage from the ER stores was sufficient to sensitize the compressed cells to apoptosis.

## Discussion

In this study, we demonstrate that volumetric compression acts as a biophysical regulator in cell state transformation. This compressive force leads to changes in the cytoskeleton, cell-ECM interaction, and proliferation. We show intracellular calcium signaling and the expression profile of intercellular calcium channels (ITPRs) were also altered, as a consequence of the adaptation process under compression conditions (Fig. 7g). The low calcium signaling in the compressed cells leads to an apoptosis-resistant phenotype.

A brief (4 hours) exposure to volumetric compression induced by hyperosmolarity has been shown to be sufficient to shift cell transcriptomic profile in lung cancer^56^. Here, using similar techniques, we found long-term (at least 5 days) adaptation to the volumetric compression induces EMT and the loss of liver specificity of hepatoblast-like liver cancer cells. The compressed cells exhibited drastic cytoskeletal reorganization, from cortical actin-rich to enhanced Rac-1 driven lamellipodia, which facilitates strong engagement between cells and the substrate. This leads to increased cell spreading, which can be also observed in other liver cancer cell types, such as, Hep3B (Fig. S7). In addition, the stronger cell-substrate engaging of the compressed HepG2 cells sustains high nuclear translocation of the transcription factor YAP (Fig. 4g), whose dysregulation was shown to be tumorigenic in multiple types of solid tumors^25,57,58^. In liver cancer patients, we found regions of packed cancer cells that were compressed against dense fibrotic capsules. These cancer cells exhibited size and density gradients near tumor boundaries (Fig. 1). We also identified regions of highly deformed peritumor normal liver cells due to the tumor expansion. Both compressed cancer cells and the deformed adjacent normal cells exhibited increased expression of YAP and higher YAP nuclear translocation (Fig. S2), supporting a link between compression and regulation of YAP. Recently, a study has shown the coactivations of YAP/TAZ in liver tumors and their peritumor cells compete with each other to determine tumor survival^59^. Whether and how mechanical compression plays a role in this YAP activation landscape in a packed liver tumor microenvironment warrants further investigation.

Recent reports proposed a novel role of colloid-induced high fluid viscosity in cell spreading and cancer cell migration triggered by calcium influx in an acute manner^60–62^. Here we show that colloid-induced hyperosmolarity also regulates cell spreading, but in a gradual way mediated by phenotypic adaptation. A distinct morphological change started to emerge on Day 3 of cell culture under compression (Fig. 3a-d). The regulations of the cytoskeleton and the actin-microtubule crosstalk are crucial in cell morphology and mechanosensing^63^. We interrogated the roles of both actin and microtubules in the compression-responding morphological changes using small molecule inhibitors (Fig. 5d-g). We found the inhibition of Rac1 or actin nucleator ARP2/3 suppressed the compression-driven spreading, indicating Rac1-facilitated actin polymerization was involved in compression-induced cell spreading. Microtubule dynamics were also important in the morphological response, as the early microtubule stabilization inhibited cell spreading under compression. Inhibiting Rho/ROCK myosin contractility did not negatively impact cell elongation, suggesting that contractility did not determine the compression-enhanced cell spreading (Fig. 5 d-g). A similar finding was also demonstrated in a study using physical compression^18^. Furthermore, suppressing Rac1 activity, but not contractility, in the compressed cells also reduced survival (Fig. 5h), which suggests the enhanced Rac1-driven cell spreading is a cell survival strategy for adaptation to compressive stress.

We further found that the compression-driven phenotype is not substrate stiffness dependent. On substrates with a physiological (∼3kPa) or pathological (∼16kPa) stiffness in the liver^46^, we also observed compression-induced cell spreading and stronger YAP nuclear translocation (Fig. S3 a,b). Previous studies have explored whether cells retain the memory of the phenotypes induced by the mechanical environments, including viscosity^62^ and substrate stiffness^64–66^. Here, we demonstrate that the cancer cells retain the memory of compression-induced enhanced cell spreading that influences cell-ECM interactions. We showed the 5-day compression-primed cells rapidly (within 24 hours) spread into much larger cells with pronounced cell protrusions and strong nuclear YAP on soft, nonelastic collagen gels (Fig. S3 c,d).

In addition to the 2D studies, we also investigated the cellular response to compression using a more physiologically relevant 3D context (Fig. S4). To recapitulate the initial stage of a tumor, we embedded HepG2 cells at the single cell level in collagen-Geltrex co-gels that mimic both fibrous ECM and basement membrane. Comparing the colony size and morphology (aspect ratio) after growth for 7 days with or without compression (Fig. S4a), we found the compressed HepG2 cells grew into smaller colonies with pronounced protrusions, whereas under the isotonic condition, the cells grew into much larger multicellular aggregates with smooth boundaries but minimum protrusions (Fig. S4b). This observation phenocopies what was seen on the 2D substrate (Fig. 2c-d).

We further found the osmotically compressed cells in 3D phenocopied mechanically compressed cells (Fig. S5). We investigated the growth patterns of HepG2 over 7 days under mechanical compression. A 20% compressive strain was applied on pure collagen ECM with a mesoporous physiological architecture that was developed previouly^54,55^. The uncompressed HepG2 cells grew into colonies against the surrounding fibrous ECM with a clear tumor boundary and rounder, uniform cell shapes (Fig. S5 b-d). When applying compression onto the encapsulating collagen gels, we found the cells grew into colonies consisting of larger and more elongated cells with distinct invasive actin protrusions that could navigate through the porous collagen structure. These observations support that compressive stress tunes the invasiveness of cancer cells and cell-ECM interactions via cytoskeleton regulations.

Studies have shown the confining cell volume leads to low intracellular calcium concentration and arrested proliferation via deactivation of TRPV4^37,38^. However, it is not well understood how intracellular calcium signaling is impacted under volumetric compression and whether there is a link between low intracellular calcium and resistance to apoptosis. Our data show that ITPR2, associated with better liver cancer prognosis based on TCGA analysis (Fig. S6), was significantly downregulated under compression. Further, ITPR3, considered a novel marker for HCC^30^ and associated with poor prognosis, was significantly upregulated in the compressed cells and enriched near the mitochondria (Fig.6 h,i). These changes in ITPR isoform expression and location may be related to the reduced calcium signaling in both the intracellular space and mitochondria upon the ATP simulation. Notably, the intracellular calcium storage in ER was unaffected (Fig. 7a,b). It is known that mitochondrial calcium uptake requires proximity with ITRPs of the ER^67,68^. Evidence also showed that reduction in ITPR3 prevents cell apoptosis via lowering the calcium influx^69^. Our observed changes in calcium responses, however, suggest the highly upregulated ITPR3 and its enhanced proximity to mitochondria in the compressed cells is insufficient to elevate calcium release, which is likely due to the reduction in ITPR2 or the low IP_3_ sensitivity of ITPR3, the lowest IP_3_ sensitivity among all three isoforms^70–72^. We found that the impaired calcium signaling de-sensitized the compressed cells to apoptosis (Fig. 2g; Fig. 7c-f), which can be circumvented by stimulated calcium mobilization by treating cells with an ER stressor. When calcium signaling is increased in this way, the compressed and non-apoptotic cells can be converted to cells that can undergo apoptosis (Fig. 7 c-f).

As microtubules are important cellular components involved in migration^73^, maintaining cell shape, and cell protrusion dynamics^74^, we also investigated microtubule dynamics as a player in the enhanced cell spreading under compression. Treatment with PTX, a microtubule stabilizer targeting mitosis to induce cancer cell death, at an early stage prevents the compressed cells from acquiring a protrusive morphology (Fig. 5 e-g). However, a microtubule destabilizer, nocodazole, showed minimal effects on the cell protrusive phenotype. With either microtubule disruptors under the compression, over 80% of the cells remained viable whereas the drug-treated, non-compressed counterparts showed significant cell death, which indicates chemoresistance induced by compression-signaling (Fig. 5i). Furthermore, in a biomimetic 3D environment, the better cell survival from PTX treatment under compression gave rise to larger tumorous colony growth, compared to the non-compressed condition (Fig. S4a,b). Along with previously reported findings^21^, we demonstrate that mechanically stressed cancer cells can lead to acquired chemoresistance and therapeutic challenges.

Despite the significant inhibition of cell proliferation under compressive stress over at least 5 days, apoptosis was not triggered. Instead, cells acquired resistance to drug-induced apoptosis (Fig. 2e-g). Under severe compression (4% PEG), there were essentially no proliferative cells between Day 5 and Day 6 (Fig. 4f). However, despite proliferation differences, in the wound healing assay, these cells exhibited a similar gap closure capability when compared to the cells maintained under isotonic conditions, and significantly higher gap closure rate than the cells under intermediate compression (2% PEG) over 8 days (Fig. 4f). This indicates different modes of wound closure - one driven by proliferation and the other driven by cell spreading and migration. The spheroid invasion assay in the stromal ECM-mimicking gels (Fig. S4d-f), as a more relevant 3D model compared to the 2D wound healing assay, better decouples bulk growth and cell invasion. Our results showed that in 3D ECMs, the hyperproliferative, non-protrusive, and non-disseminating HepG2 cells are converted into a non-proliferative, invasive phenotype under compression.

Our study demonstrates adaptation modes acquired by liver cancer cells under volumetric compression, notably cytoskeletal changes (mediated by Rac1) and impaired calcium signaling. The adapted cell states are anti-apoptotic, drug-resistant, and more invasive - all of which drive worse prognosis. Notably, elevating intracellular calcium baseline re-sensitizes the compressed cells to apoptosis. Our findings highlight the interplay between the mechanical regulation (particularly volumetric compression) and liver cancer progression and offer new potential treatment paradigms that directly target and counter compression-adapted tumor cell subpopulations.

## Materials and Methods

Methods and materials used for this study are described in SI Appendix.

### Statistics

The number of biological replicates and technical replicates, as well as the methods of statistical comparisons, were stated in figure legends. The difference is considered significant when the *p*-value is less than 0.05. Comparisons with no significant difference are labeled as “ns”. The statistical comparisons were performed using GraphPad Prism 9 (GraphPad). All bar plots represent mean ± standard deviation (SD). Boxes, lines in the boxes, and the whiskers in the box plots represent 25th to 75th percentiles, median, minimum, and maximum. Lines in the violin plots represent the median and 25th to 75th percentiles.

## Acknowledgements

We thank Dr. Michael H. Nathanson from Yale Liver Center for reagents and technical assistance. We also thank Dr. Xuchen Zhang from Yale Surgical Pathology for the histology samples of liver cancer patients and his valuable inputs. We acknowledge Saltzman Lab for the access to their plate reader. We thank the Liver Center at UFMG (Brazil) for valuable discussions. We acknowledge the technical support from Yale Center for Genome Analysis (YCGA). This project was also supported in part by the Yale Liver Center award NIH P30 DK034989 Cellular and Molecular Physiology Core Facility. M.M. discloses support for this study from Yale Liver Center Pilot Project under award NIH P30 DK034989, and NIH R35GM142875. R.Y.N. discloses support for this study from NIH grant T32EB019941. M.F.L. disclosed support for this study from CAPES Print, CNPq, FAPEMIG (Brazil).

