## Supplementary material for "Adaptation to volumetric compression drives hepatoblastoma cells to an apoptosis-resistant and invasive phenotype": SI Appendix

### Methods

**Mechanical signature analysis on HCC histological samples.** HCC cases were selected from the Yale New Haven Hospital pathology database. The pathology slides were evaluated for background liver disease and the presence or absence of cirrhosis. The images of the pathology slides were acquired using an imaging service from HistoWiz (https://home.histowiz.com/). Nuclei detection was adapted from the StarDist segmentation algorithm[75,76]. Nuclear area and nuclear density within a 30 μm radius of each nucleus are the two metrics to determine the signature. Metrics for each nucleus were mapped onto the original histology slides with a colormap representing the magnitude of the measurements. Circles in the overlayed images represent the center of mass of each segmented nucleus. The tumor capsule and the boundary between the tumor and the adjacent normal were manually traced, and the minimum distance between each nucleus' center of mass was calculated. Scatter plots were then generated to visualize each metric with respect to distance from the selected tumor boundaries. All scripts are available upon reasonable request.

**Cell maintenance and culture in osmotic compression medium.** Hepatoblast-like HCC cell line HepG2 was cultured in DMEM with 4.5 g/L D-Glucose and L-Glutamine (Gibco), 10% FBS (Gibco), and 1% Pen Strep (Gibco). The cells were maintained in a humid incubator at 37°C with routine medium change every 2-3 days. To prepare two different compression media, filter-sterilized PEG300 was added at concentrations of 2% and 4% (vol/vol). To match the slight dilution of the serum in the compression medium by the addition of PEG300, our isotonic "control" medium was the regular culture medium added with 4%(vol/vol) 1×PBS. To ensure the full incorporation of PEG into the medium, we prepared the PEG condition media a week before using them. For the 2D compression experiments (illustrated in Fig. 2a), we released the cells from culture flasks by 5-min 0.25% Trypsin-EDTA (Gibco) treatment and removed large cell clumps with a 40 μm cell strainer. The cells were seeded and allowed to attach to tissue culture well plates or glass 8-chamber slides (Cellvis) for 1 day in the regular culture medium. We then switched the medium to isotonic, 2% PEG, or 4% PEG medium and continued culturing the cells for 5 to 7 days.

**RNA-seq and analysis.** Biological replicates of HepG2 cells cultured in isotonic and 4% PEG medium for 5 days were collected for bulk RNA sequencing. Total RNA was extracted with an RNeasy mini kit (QIAGEN), following the manufacturer's protocol. The library was prepared and sequenced on the Illumina HiSeq platform (paired-end 100bp reads per sample) by Yale Center for Genome Analysis (YCGA). The raw reads were aligned by a read aligner STAR[77] to the human GRCh38 release-86 transcriptome (https:// www.ensembl.org/). Data normalization and differential expression analysis were conducted with the R-package DESEQ2 (https://bioconductor.org/packages/release/bioc/html/DESeq2.html) on RStudio. We considered genes differentially expressed if the adjusted P value was less than 0.05. For pathway and process, we conducted both gene set enrichment analysis (GSEA) (https://www.gsea-msigdb.org/gsea/index.jsp) and analysis on METACORE (https://portal.genego.com/). Volcano plots were also made to visualize differentially expressed genes using an R-package (https://bioconductor.org/packages/release/bioc/html/EnhancedVolcano.html).

**Reagents and drugs.** The primary antibodies used in this study are: anti-YAP (sc-101199, Santa Cruz, 1:200), anti-β catenin (D10A8, Cell Signaling Technology, 1:200), anti-ITPR1 (Alomone labs, ACC-019), anti-ITPR2 (Santa Cruz Biotechnology, sc-398434), anti-ITPR3 (BD Transduction Laboratories, 610313), anti-cleaved caspase 3 (Cell Signaling Technology, #9661), anti-TOM20 (D8T4N, Cell Signaling Technology), anti-GAPDH (Thermo Fisher Scientific, AM4300). The secondary antibodies used in this study are: Alexa Fluor 647 Goat Anti-Mouse IgG antibody (Invitrogen) and Alexa Fluor 488 Goat Anti-Rabbit IgG antibody (Invitrogen). We use Hoechst 33342 (Invitrogen, H3570, 1:2000) to stain the cell nuclei, and Rhodamine Phalloidin (Abcam, ab235138, 1:1000) or Alexa Fluor 488 Phalloidin (Invitrogen, A12379, 1:500) to stain for F-actin. Small molecule drugs used in this study are: ROCK inhibitor Y27632 (Cayman, 10μM), Rac1 inhibitor NSC23766 (Tocris, 100μM), ARP2/3 inhibitor CK666 (Cayman, 100μM) microtubule depolymerizer Nocodazole (Sellekchem, 100nM), microtubule stabilizer Paclitaxel (Sellekchem, 100nM), apoptosis inducer Staurosporine (Sigma, 5μM), sarco/endoplasmic reticulum Calcium ATPase (SERCA) inhibitor Thapsigargin (Sigma, 50nM or 1μM).

**Immunofluorescence staining.** Cells were washed with PEG-conditioned PBS and then fixed with PEG-conditioned 4% paraformaldehyde (PFA) (Santa Cruz) for 15 min at room temperature (RT). PFA was also accordingly conditioned to minimize osmolarity shock before a full fixation. After washing with PBS three times, the cells were permeabilized with 0.3% Triton X-100 in PBS for 15 min, washed with PBS, and blocked in 1% bovine serum albumin (BSA) in PBS for 1 hour at RT. After blocking, samples were incubated with the primary antibodies in PBS overnight at 4°C. Followed by a thorough wash in PBS for 1 hour, the samples were stained with the secondary antibodies and rhodamine-phalloidin for F-actin in the dark for 1 hour at RT. After a thorough wash in PBS for 1 hour, the cells were then stained with nucleus dye Hoechst before the imaging.

**RhoA and Rac1 activation assays.** Colorimetric-based GLISA assays (Cytoskeleton, Inc.) were used to quantify RhoA and Rac1 activity. The manufacturer's manuals were strictly followed. In brief, cells cultured in the 6-well plates in isotonic, 2% PEG, and 4% PEG conditions were quickly washed with ice-cold PBS and lysed with cold lysing buffer. The protein solution was clarified by 1 min high-speed centrifuging (10,000 RPM) at 4°C, aliquoted, snap-frozen in liquid nitrogen, and stored at -80°C. The whole procedure was finished within 10 min to prevent excessive hydrolysis of the active G-proteins. A small aliquot of cell lysate was used for total protein quantification with a "BCA Protein Assay Kit" (23225, Thermo Scientific). The same quantity (60 μL) of total protein from each condition at the concentration of 0.5 mg/mL was loaded in duplicate or triplicate wells of the GLISA plate. The procedures of protein binding, antigen-presenting, conjugation of primary and secondary antibodies, and signal detection followed the manufacturer's manual. The signal in each well was detected by a microplate spectrophotometer (Molecular Devices) at the absorbance of 490 nm.

**Transfection of LifeAct-labeled HepG2.** HepG2 cells (~$2\times10^4$ cells) in a well of a 24-well plate were incubated in 500 μL complete RPMI medium with 2μL (2 multiplicity of infection) rLVUbi-LifeAct-TagRFP lentiviral vector (Ibidi) and 8 μg/mL polybrene (Sigma) for 16 hours. The medium was then replaced with fresh medium and the cells were allowed to expand for 2 days. Stably transfected (RFP positive) cells were selected with 2 μg/mL puromycin (Gibco) in the culture medium.

**Tumor growth in the 3D niche.** HepG2 spheroids were made by aggregating 1,000 cells in 100uL medium per well in an ultra-low attachment 96 well round bottom plate (CORNING) for 4 days. The spheroids were collected with wide-bore pipette tips and stored in the medium before the experiments. Random embedding of single cells and small cell clusters was also used in the 3D studies. Basement membrane (BM)-collagen co-gel solution was prepared by mixing equal volumes of ice-cold reduced growth factor basement membrane matrix Geltrex (A1413202, Thermo Fisher) and ice-cold neutralized 4mg/mL rat tail collagen to achieve final concentrations of 50% and 2mg/mL for BM and collagen, respectively. The 4mg/mL collagen was diluted with prechilled cell culture water and 10x PBS from a ~10mg/mL acidic collagen stock (354249, Corning), and the pH was then adjusted by 0.5M NaOH on ice. For the spheroid studies, 4-5 spheroids were incorporated in 50uL co-gel solution per well in glass bottom 12-well plates (Cellvis). For the loose cell embedding, ~5000 cells were incorporated in ~25uL co-gel solution per well in 8-chambered glass slides (Cellvis). The glass bottom plates or slides were coated with dopamine-HCl (Acros Organics) to promote gel anchorage[26]. The co-gel droplets formed domes and were allowed to solidify for 40 min at 37°C, followed by the addition of isotonic or PEG-conditioned media with or without drug treatments. The spheroids or loosely embedded cells were allowed to grow for 7 days, with a medium change every 2-3 days.

**Calcium imaging.** For intracellular calcium measurements, HepG2 cells were incubated with 6 μM Fluo-4/AM (Invitrogen, F14201) for 30 minutes at 37°C, transferred to a perfusion chamber on the stage of a Zeiss LSM 710 confocal microscope (Carl Zeiss, Inc., Thornwood, NY), perfused with PEG300-conditioned HEPES-buffered solution (NaCl, 130 mmol/L; KCl, 5 mmol/L; $CaCl_2$, 1.25 mmol/L; $KH_2PO_4$, 1.2 mmol/L; $MgSO_4$, 1 mmol/L; HEPES, 19.7mmol/L; glucose, 5 mmol/L; pH 7.4) while stimulated with 1 μM of adenosine triphosphate (ATP). For ER calcium measurements, cells was perfused with PEG300-conditioned HEPES-buffered solution while stimulated with 1 μM of thapsigargin. Calcium signaling was monitored by exciting at 488 nm while collecting emitted light above 505 nm. Fluo-4 fluorescence was monitored using a 40X, 1.2 NA objective lens, and images were collected at a rate of 1-5 frames/second. The mitochondria calcium was monitored using adenovirus-encoded mitochondrial calcium indicator (ad-mito-R-GECO)[78]. Changes in fluorescence F were normalized by the initial fluorescence ($F_0$) and were expressed as ($F/F_0$) x 100%. Normalized amplitudes of ATP-induced calcium signals were extracted with ImageJ software (NIH, Bethesda, MD) and plotted as described[79,80].

**Mitochondria extraction**

HepG2 cells were cultured in isotonic, or 4% PEG medium in 6-well plates for 5 days and then collected using TrypLE™ Express Enzyme (Gibco, 12605010). Then, following the manufacturer's protocol for the Mitochondria Isolation Kit for Cultured Cells (Thermo Fisher Scientific, 89874), cytosol and mitochondrial fractions were obtained by multiple centrifugations after gentle disruption of the collected cells. Each fraction was used in the following experiments in exactly the same way as the immunoblot technique as described below.

**Cell viability assay.** Propidium iodide (PI) -based Cell Viability Kit (R37610, Invitrogen) was used to stain dead cells. Cells were cultured in 8-chambered cover glass (Cellvis) under different compression conditions with drug treatments. On Day 5, the ratio of 2 drops/mL of PI was added in the chambers supplemented with a nucleus stain Hoechst (dilution: 1:2000) as the counterstain. After 30-min incubation at 37°C, the cells were imaged at 360/460 nm and 535/617 nm on a confocal microscope to detect the total cells and the dead cells. Viability was calculated as a percentage of PI stain-negative cells out of the total cells.

**Cell apoptosis imaging-based assay.** CellEvent Caspase-3/7 Green Detection Reagent (C10423, Invitrogen) was used to label caspase-3/7 positive cells as a marker for cell apoptosis. The drug-treated cells under different compression conditions were incubated with CellEvent (dilution: 1:1000) and the counterstain was Hoechst (dilution: 1:2000) at 37°C for 1 hour. The cells were imaged at 360/460 nm and 502/530 nm to detect the total cells and the Capase-3/7 positive cells. The apoptotic population was calculated as a percentage of caspase-3/7 positive cells out of the total cells.

**Cell proliferation (EdU) assay.** Cell proliferation was assessed using EdU Cell Proliferation Kit, Alexa Fluor 555 dye (C10638, Thermo). HepG2 cells were seeded on the 8-chambered glass slides and cultured in isotonic or PEG-conditioned media for 5 days. On Day 5, the cells were switched in the same type of culture media that were supplemented with 1:1000 EdU for 12 hours. Proliferative cells over the 12 hours were tagged with EdU. The cells were then fixed and conjugated with the fluorescent dye, following the manufacturer's manual. Hoechst (1:2000) was used as the counterstain. The cells were imaged at 360/460 nm and 555/565 nm to detect the total nuclei and the EdU positive nuclei. The proliferation was calculated as a percentage of EdU positive cells out of the total cells.

**Immunoblotting.** HepG2 cells were cultured in isotonic, 2% or 4% PEG medium in 6-well plates for 5 days and then collected. When experiments were performed with staurosporine or thapsigargin (50nM), it was administered on day 5 and collected 12 hours later (day 6). All reagents used in the following procedures were purchased from Thermo Fisher Scientific, except as noted. Cell samples were lysed in RIPA Buffer (#89901) plus protease and phosphatase Inhibitor cocktail (#78440, Thermo Fisher). Bradford assay (#23200, Thermo Fisher) was used to determine protein concentrations, and loading samples were diluted with sample buffers (NP0007 & NP0004, Thermo Fisher), then boiled for 10 minutes at 70 °C according to the manufacturer's protocol. These samples were loaded onto NuPAGE™ Protein Gels. For the detection of low molecular weight proteins, MES Buffer (NP0002, Thermo Fisher) was used instead of MOPS Buffer (NP0001, Thermo Fisher). Proteins were transferred to the nitrocellulose membrane (Bio-Rad #1620145). Membranes were blocked with 5% milk (Omniblok™ , AmericanBIO, AB10109) for 30 min at room temperature and reacted with primary antibody (1:1500) overnight at 4°C. Secondary antibodies, anti-rabbit (Invitrogen, #A 27036) or anti-mouse (Cytiva, NA931) were used at 1:3000 for 1 hour at room temperature. Pierce™ ECL Western Blotting Substrate (#32106, Thermo Fisher) was used for detection. Intensities of bands on immunoblots were quantified by densitometry analysis by Image J software (NIH, Bethesda, MD) and normalized by GAPDH.

**Reverse transcription-quantitative polymerase chain reaction (RT-qPCR).** HepG2 cells were collected after 1, 3, and 5 days of culture in isotonic, 2% or 4% PEG media in 6-well plates. Total RNA was extracted according to the manufacturer's protocol for the RNeasy mini kit (QIAGEN). 500 nM of each sample RNA was used to obtain cDNA following the manufacturer's protocol for the iScript™ cDNA Synthesis Kit, (Bio-Rad, #1708891). RT-qPCR was performed using QuantStudio 6 and 7 Flex Real-Time PCR Systems (Thermo) following the manufacturer's protocol for FastStart Universal Probe Master (Merk, #4913949001). Primers used were ITPR1 (hs00181881_m1), ITPR2(hs00181916_m1), ITPR3(hs01573539_m1), 18s(hs03003631_g1) purchased from Thermo Fisher Scientific.

**Fabrication of soft fibrous collagen substrate**. A 25-µL neutralized collagen droplet was gently dispensed onto dopamine-HCl coated glass surface to form a dome in a glass bottom 12-well plate (Cellvis). The droplet was quickly flattened and sandwiched between a top BSA-treatment round coverslip (8mm in diameter) and the glass bottom surface. After gelation at 37°C in a humid incubator for 1 hour, the top coverslip was removed by gentle washing with PBS. The collagen formed a thin (~250-300 µm thickness) 3D cushion with an even surface. To visualize collagen fibers, the collagen gels were stained in Alexa Fluor 647 NHS Ester dye (12-20 µg/mL) (Thermo Fisher) in PBS at 37°C for 1 hour and were thoroughly washed with sterile PBS 5 times before seeding cells.

**Fabrication of PA substrates with physiological stiffnesses.** We followed the protocol described previously to fabricate the physiological (~3 kPa) and pathological (~16 kPa) PA substrates[81]. In brief, clean glass surfaces of 35-mm glass-bottom dishes were silanized with (3-aminopropyl)trimethoxysilane (Thermo) for 3 min followed by 0.5% glutaraldehyde (Polysicences, Inc.) treatment for 30 min. The dishes were thoroughly washed with deionized water and dried in a 60 °C oven. The 3-kPa PA gel was made by mixing 100 µL 40% acrylamide (AA) (Sigma), 112.5 µL 2% bis-acrylamide (bis-AA) (Sigma), and 787.5 µL sterile water. The 16-kPa gel was made by mixing 250 µL 40% acrylamide (AA) (Sigma), 75 µL 2% bis-acrylamide (bis-AA) (Sigma), and 675 µL sterile water. Polymerization of the PA gels was initiated by introducing 1 µL N,N,N′, N′-tetramethylethylenediamine (TEMED) (MP Biomedicals, LLC) and 10 µL 10% ammonium persulfate (APS) (Sigma). 15 µL of the final mix was sandwiched by the silanized glass surface and a top 12mm×12mm square coverslip treated by dichlorodymethyl silane (Sigma). The gel mix was allowed to polymerize for 30 min before the coverslip was removed. The PA substrates were then UV sterilized cycle in sterile PBS for 30 min. The PA substrates were later functionalized with 0.2mg/mL Sulfo-SANPAH (Thermo Fisher) followed by incubation in 50 µg/mL collagen I solution in 0.2N acidic acid at 4°C overnight. The substrates were washed three times with sterile PBS before seeding cells.

**Physical compression of HepG2 cells in 3D collagen ECM.** A UV-sterilized silicone sheet of 500 mm thickness was punched with a 4 mm radius punch biopsy to create a stencil for a collagen gel disk with a radius of 4 mm and a height of 500 mm. HepG2 cells were encapsulated in a collagen gel with mesoscopic architectures we termed “islands”. The collagen island gels were made as previously described[54]. Briefly, 350 µl of 2mg/mL type I collagen was neutralized to a pH of 7.4 and allowed to sit at room temperature for 6.5 minutes. Gels were then sheared with a 200µL pipette set to 175 µl for 3 minutes with a 5-second period. Cells were then mixed into the gel at a final concentration of 15,000 cells/mL. A 26 µL droplet of collagen solution with cells was plated on a polydopamine-coated glass surface within the sterilized stencil and a 0.3% BSA-coated glass disk was placed on top to achieve a flat top face. After 45-min gelation at 37 °C, the glass disk and silicone stencil were carefully removed to leave an intact cylindrical collagen disk gel. Thinner spacers (400 mm in height) were placed on either side of the gel with a glass slide placed on top to physically compress the gel by 20% in the z-direction. A PDMS block was placed on top of the glass slide to keep it in place. Cells were cultured in the complete DMEM medium for 7 days, fixed, and stained for nuclei (Hoechst) and actin (phalloidin). Collagen was imaged using confocal reflectance mode.

**Imaging analysis.** Cell shapes were segmented with ImageJ or manually traced based on the fluorescent imaging of actin or bright field. YAP subcellular localization was determined by the ratio of YAP mean intensity in the nuclei and cytoplasm following the quantifying method published previously[82]. Masks of the nucleus or whole cell shape were manually traced or segmented with ImageJ.

**Survival analysis from TCGA database.** Survival analysis (60 months) as a function of the genes (ITPR1, ITPR2, ITPR3) from 364 liver cancer patients (LIHC) from The Cancer Genome Atlas Program
(TCGA) (https://www.cancer.gov/about-nci/organization/ccg/research/structural-genomics/tcga) was conducted using a peer-reviewed online tool Kaplan–Meier Plotter (https://kmplot.com/analysis/)[83]. The cutoff value for the "High" and "Low" groups in the survival analysis was determined as the best-performing value among all possible cutoff values.

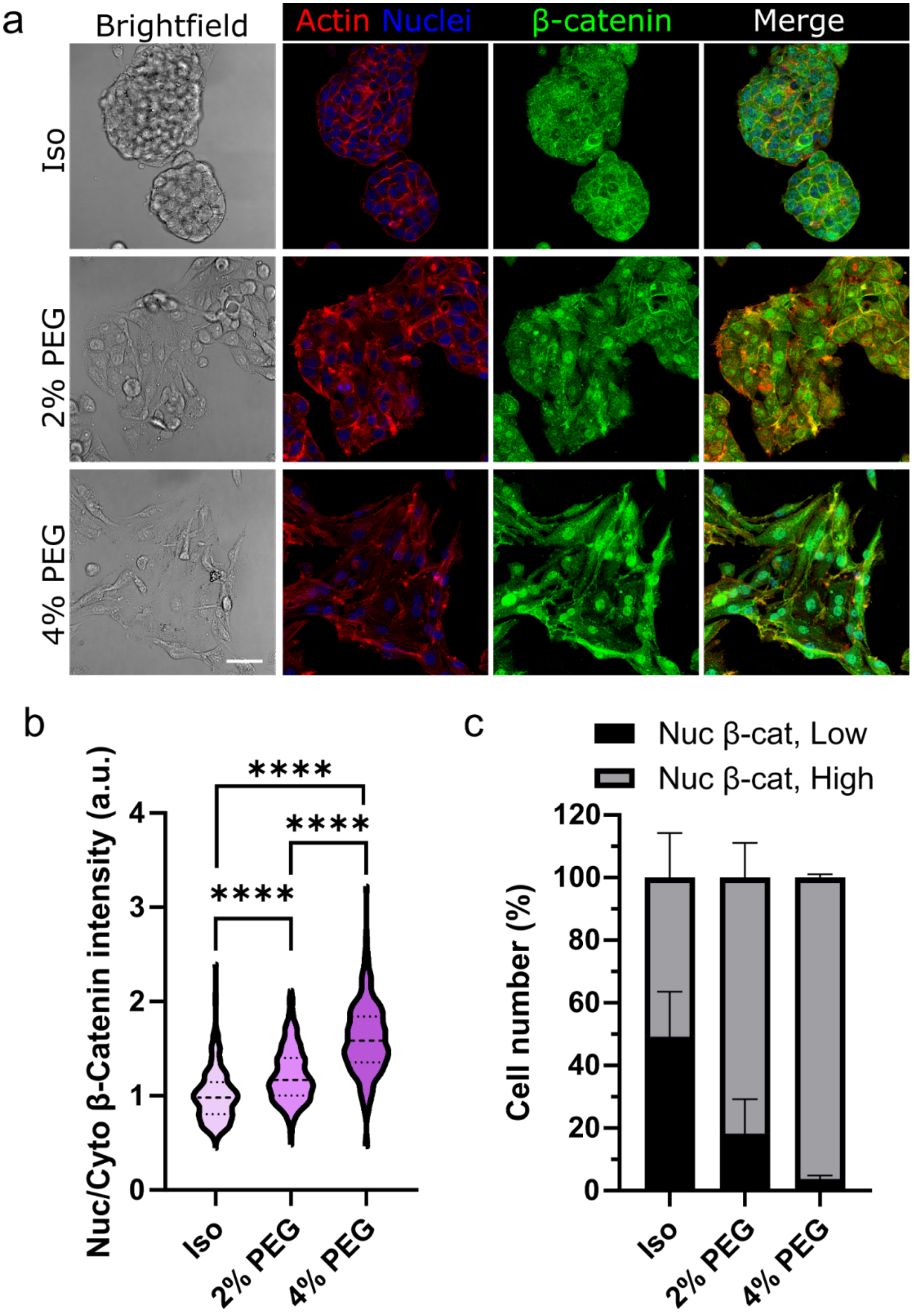

a
Brightfield
Actin Nuclei
β-catenin
Merge
Iso
2% PEG
4% PEG
b
Nuc/Cyto β-Catenin intensity (a.u.)
****
****
****
0
1
2
3
4
Iso
2% PEG
4% PEG
c
Nuc β-cat, Low
Nuc β-cat, High
Cell number (%)
0
20
40
60
80
100
120
Iso
2% PEG
4% PEG

**Figure S1. Volumetric compression drove the nuclear translocation of β-catenin.** (a) Immunofluorescent staining showing cell shape, actin organization, and β-catenin subcellular location. Scale bar: 50μm. The extent of β-catenin nuclear translocation was determined by (b) the fluorescent intensity ratio of β-catenin in nuclei and cytoplasm, and (c) the percentage of cells with high nuclear β-catenin vs. low nuclear β-catenin within instinct colonies. For (b, c) n=242-350 cells from N=3-4 individual colonies. The threshold of "high nuclear β-catenin" is "Nuc/Cyto β-catenin intensity > 1". Multiple comparisons in (b) were made using one-way ANOVA with Tukey post hoc (****$p$<0.0001).

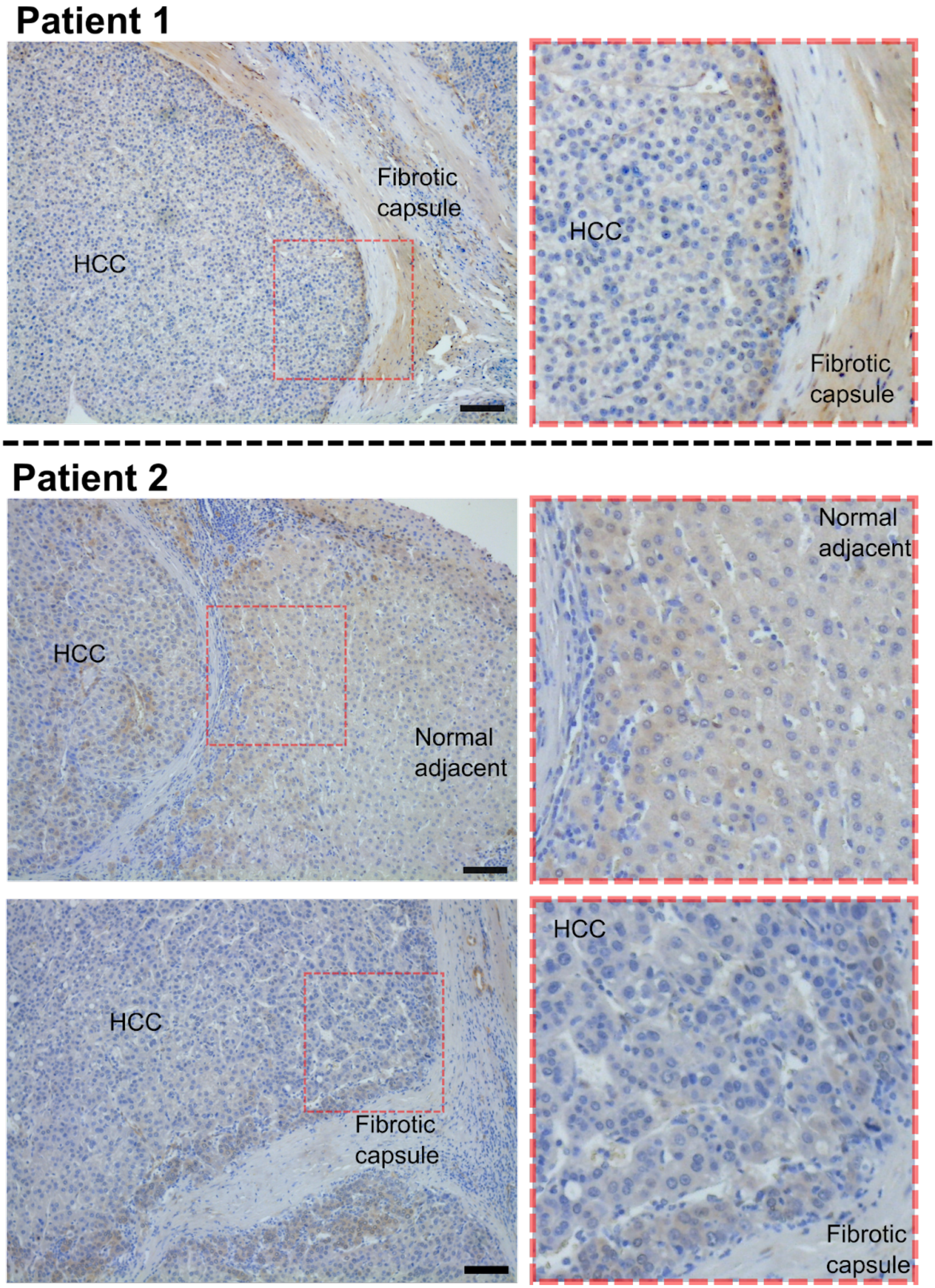
Patient 1
Fibrotic capsule
HCC
HCC
Fibrotic capsule
Patient 2
HCC
Normal adjacent
Normal adjacent
HCC
Fibrotic capsule
HCC
Fibrotic capsule

**Figure S2. Spatial distribution of YAP expression in HCC patients.** Denser regions of HCC near fibrotic stroma exhibited higher expression of both total YAP and nuclear YAP.

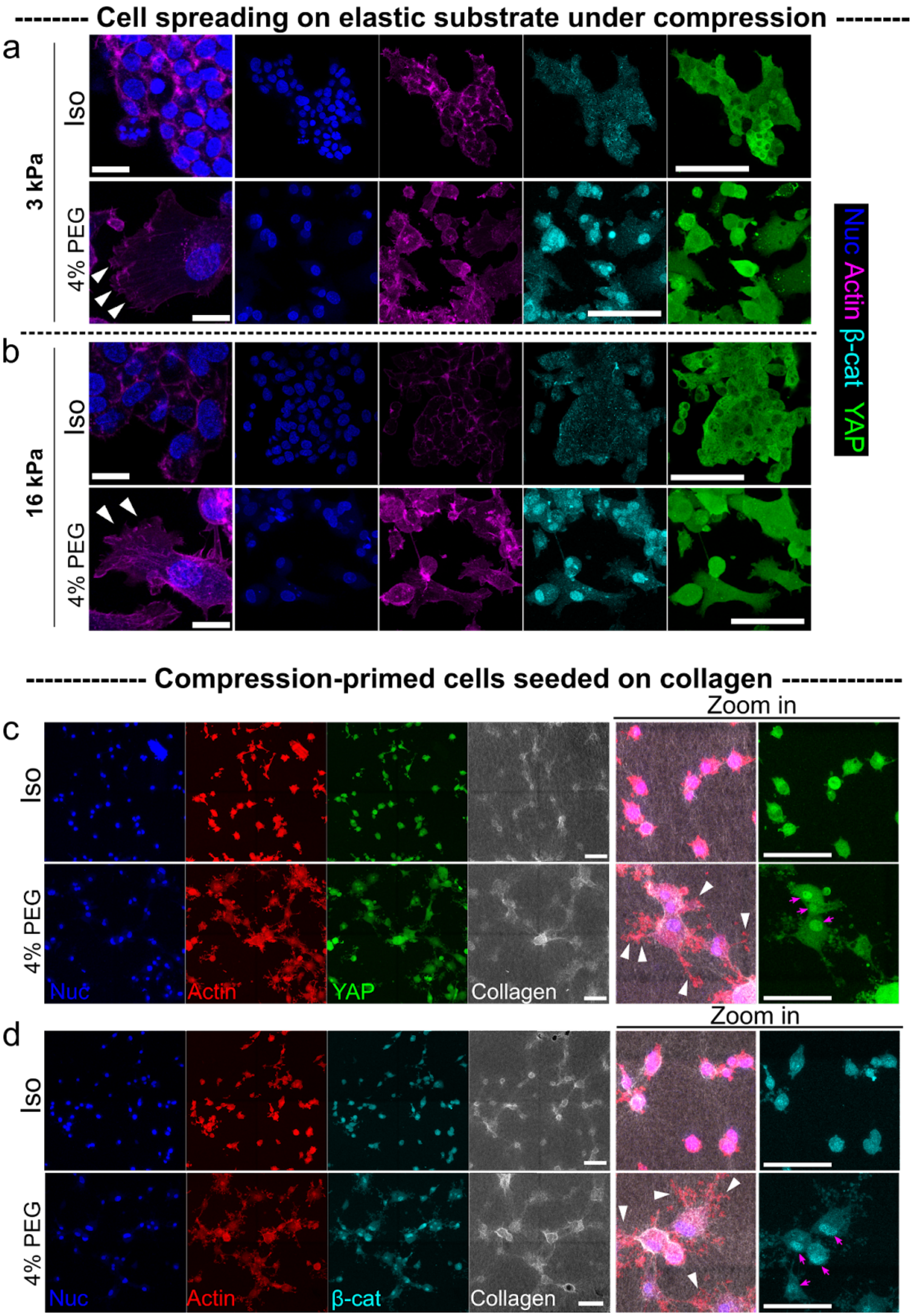

-------- Cell spreading on elastic substrate under compression --------
a
3 kPa
Iso
4% PEG
b
16 kPa
Iso
4% PEG
Nuc Actin β-cat YAP
------------- Compression-primed cells seeded on collagen -------------
c
Zoom in
Iso
4% PEG
Nuc
Actin
YAP
Collagen
d
Zoom in
Iso
4% PEG
Nuc
Actin
β-cat
Collagen

**Figure S3. Volumetric compression enhanced cell spreading and nuclear translocations of YAP and β-catenin on physiologically relevant substrates.** We cultured HepG2 cells on the elastic PA-gel substrates with (a) a physiological stiffness (3kPa) and (b) a pathological (cirrhosis) stiffness (16kPa) in the isotonic condition and the 4% PEG condition. On both substrates, HepG2 cells grew into cell-packed colonies mainly with cortical actin in the isotonic condition for 5 days. In comparison, the compressed cells exhibited distinct lamellipodia-like protrusions (indicated by the arrowheads), large cell spreading areas, and stronger nuclear translocations of both β-catenin and YAP. In (c) and (d), We also found cells compressed on the 2D rigid substrate for 5 days, which we termed as "compression-primed cells", sustained the phenotypes of the enhanced cell spreading area and pronounced protrusion formation (indicated by the arrowheads), 24 hours after being detached and re-seeded on the fibrous collagen (2mg/mL) gel surface in the same PEG-conditioned medium. Compared to the non-compressed cells in the isotonic condition, the compression-primed cells also showed higher nuclear translocation of both YAP and β-catenin (indicated by the magenta arrows). The collagen was fluorescently labeled to show the remodeling of the ECM by cellular force. Scale bars in the first columns (the "zoom in" images) of (a) and (b) are 20 μm. The other scale bars are 100μm.

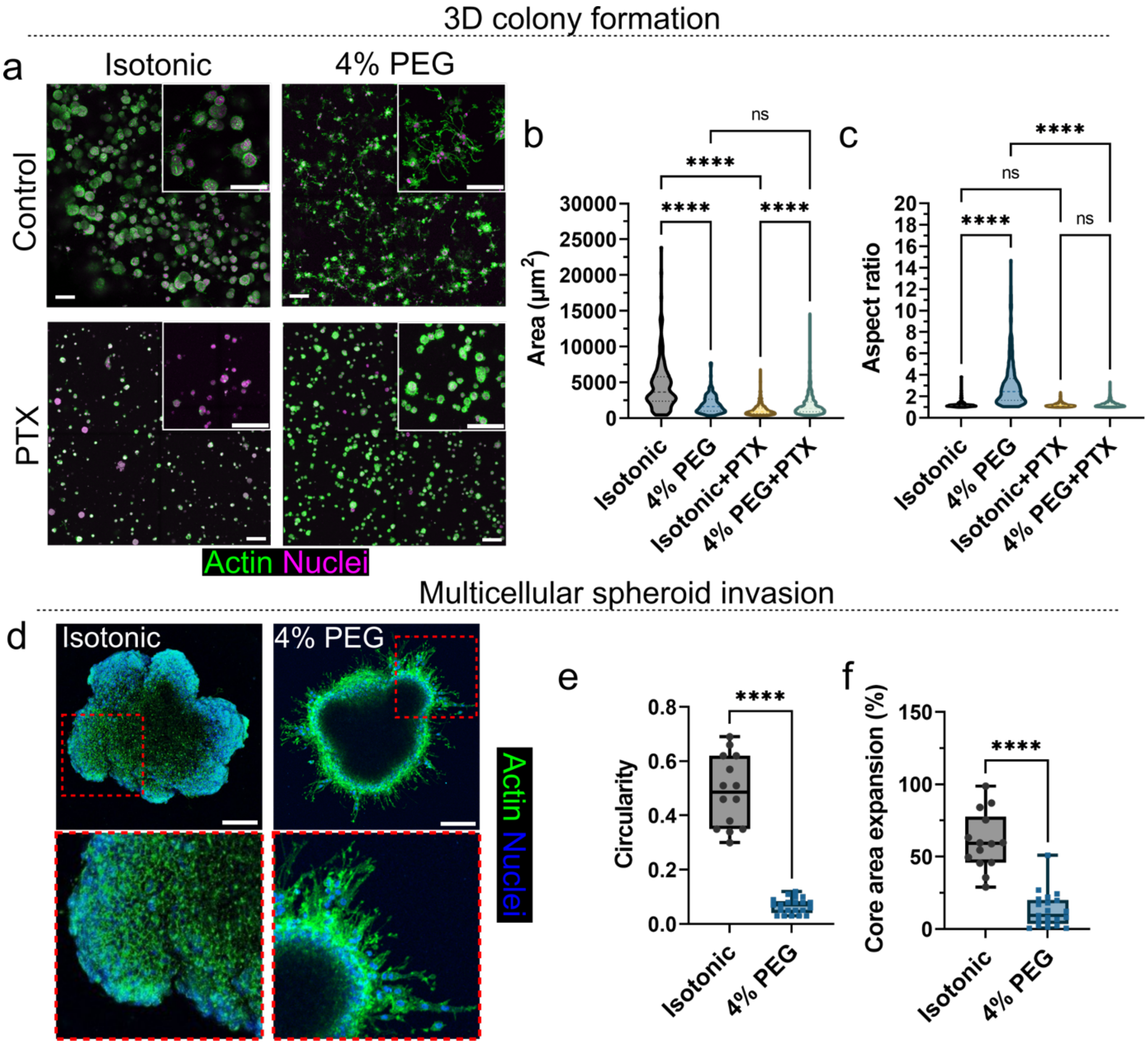


**Figure S4. Compression leads to an invasive, non-proliferative liver cancer phenotype.** (a) Seven-day growth of single cells embedded in collagen-BM co-gels under compression and drug treatments. 4% PEG compression suppresses multicellular colony formation but promotes cell protrusions to breach the BM barrier. When treated with PTX, the colonies grew bigger under compression than those in the isotonic condition. Cell/colony morphology is characterized by (b) aspect ratio and (c) projected area. n= 185, 248, 418, 872 aggregates for Isotonic, 4% PEG, Isotonic+PTX, and 4% PEG+PTX respectively from N= 2 independent experiments. As observed in the 2D experiments, treatment with PTX also diminished the compression-induced protrusion formation in the 3D context. Moreover, the compression-induced desensitizing to PTX rendered a better cell survival, leading to bigger colony sizes in 3D under compression. (d) Multicellular tumor spheroids embedded in the co-gels exhibited a distinct difference in cell-ECM interactions under compression. The tumors in the isotonic condition exhibited expansion against the ECM without invasive cells observed. The compressed tumors showed distinct individual cells breaching the ECM encapsulation and migrating out of the tumor core onto the gel-substrate interface. (e) Tumor expansion was characterized by the core area change compared to the tumor area on Day 0, showing the spheroids encapsulated in the co-gel in the isotonic condition expanded by ~60% in footprint with minimum cell processes observed in the ECM, recapitulating a rapid, bulk expansion of HCC tumors against the stroma. The volumetric compression, in contrast, significantly reduced the tumor expansion, yet led to cell protrusion formation, the breach of ECM, and the subsequent invasion as multicellular strands or single cells. (f) Cell invasion was characterized by the circularity of the Day 7 tumor shape contour. The overall outward invasion was dramatically increased highlighted by the reduction of the tumor circularity. n= 14 and 21 individual spheroids for Isotonic and 4% PEG respectively from N= 3 independent experiments. A two-tailed student's t-test was performed for (e) and (f).

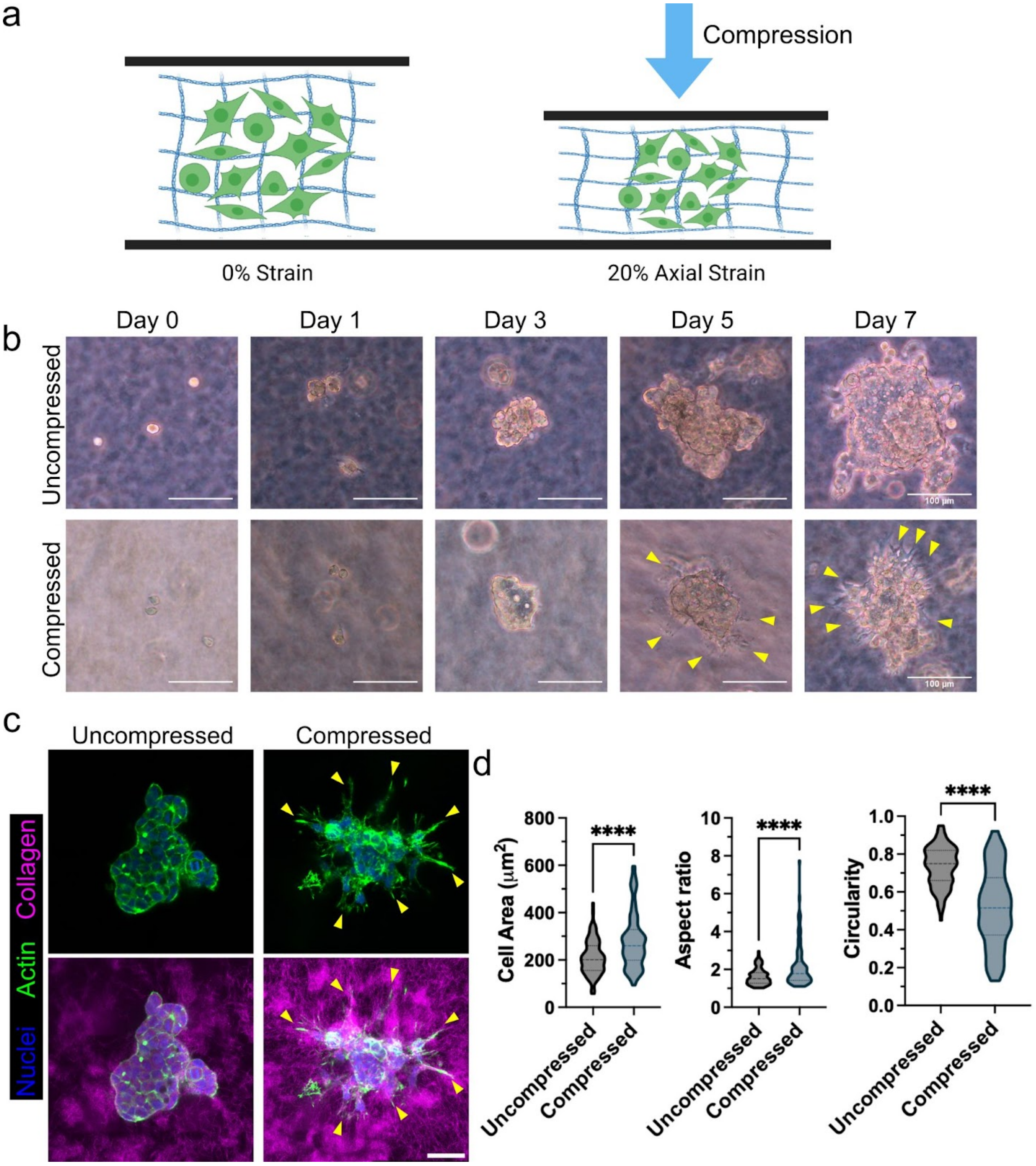


**Figure S5. Mechanical compression drives protrusive phenotype and stronger cell-ECM interaction.** (a) Schematic of the experimental setup for mechanically compressing HepG2 in a 3D collagen niche. (b) Representative images of HepG2 cells growing in collagen ECM with or without mechanical compression on Day 0, 1, 3, 5, 7. On Day 5 and Day 7, distinct protrusive cells were observed. (c) Fluorescent images comparing cytoskeleton arrangements of HepG2 colonies in collagen ECM with vs. without mechanical compression. Yellow arrows indicate cell protrusions that exhibit strong interactions with the surrounding fibrous collagen. (d)

Quantification of the morphology (cell area, aspect ratio, and circularity) of individual cells in the multicellular colonies with or without mechanical compression. In the compressed environment, the cells became larger and more elongated. n=111 cells for the "uncompressed"condition, and 104 cells for the "compressed" condition, measured from four multicellular colonies. A two-tailed student's t-test was performed in (d) (****$p<0.0001$). Schematic in (a) was made using BioRender.

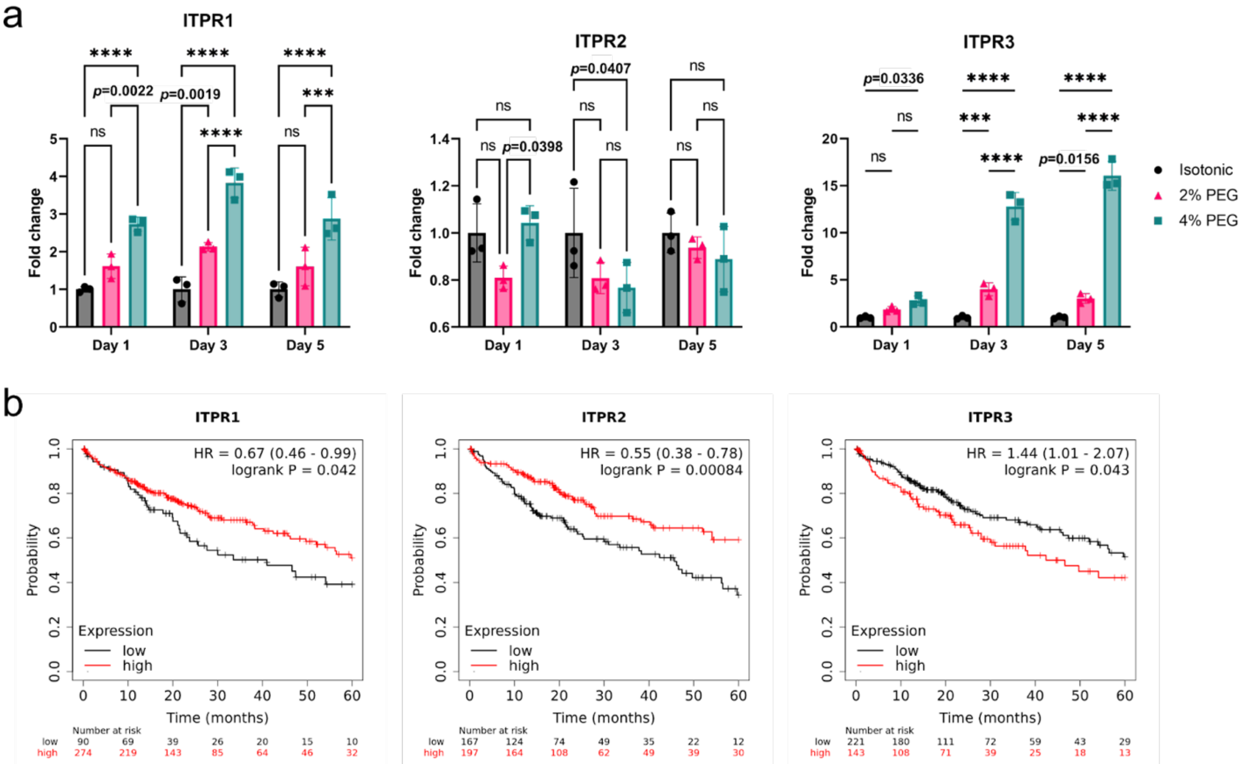


**Figure S6. Gene expressions of ITPRs in HepG2 under compression and TCGA survival analysis on liver cancer patients.** (a) qPCR reveals dysregulated InsP3 receptors (ITPRs) under volumetric compression as a function of time. N=3 biological replicates. Two-way ANOVA with Tukey post hoc was used for the statistical analysis (***$p$<0.001, ****$p$<0.0001). (b) TCGA analysis suggests a linkage between ITPRs and liver cancer prognosis. Upregulations of ITPR1 and ITPR2 show a good prognosis, and overexpression of ITPR3 is associated with a poor prognosis.

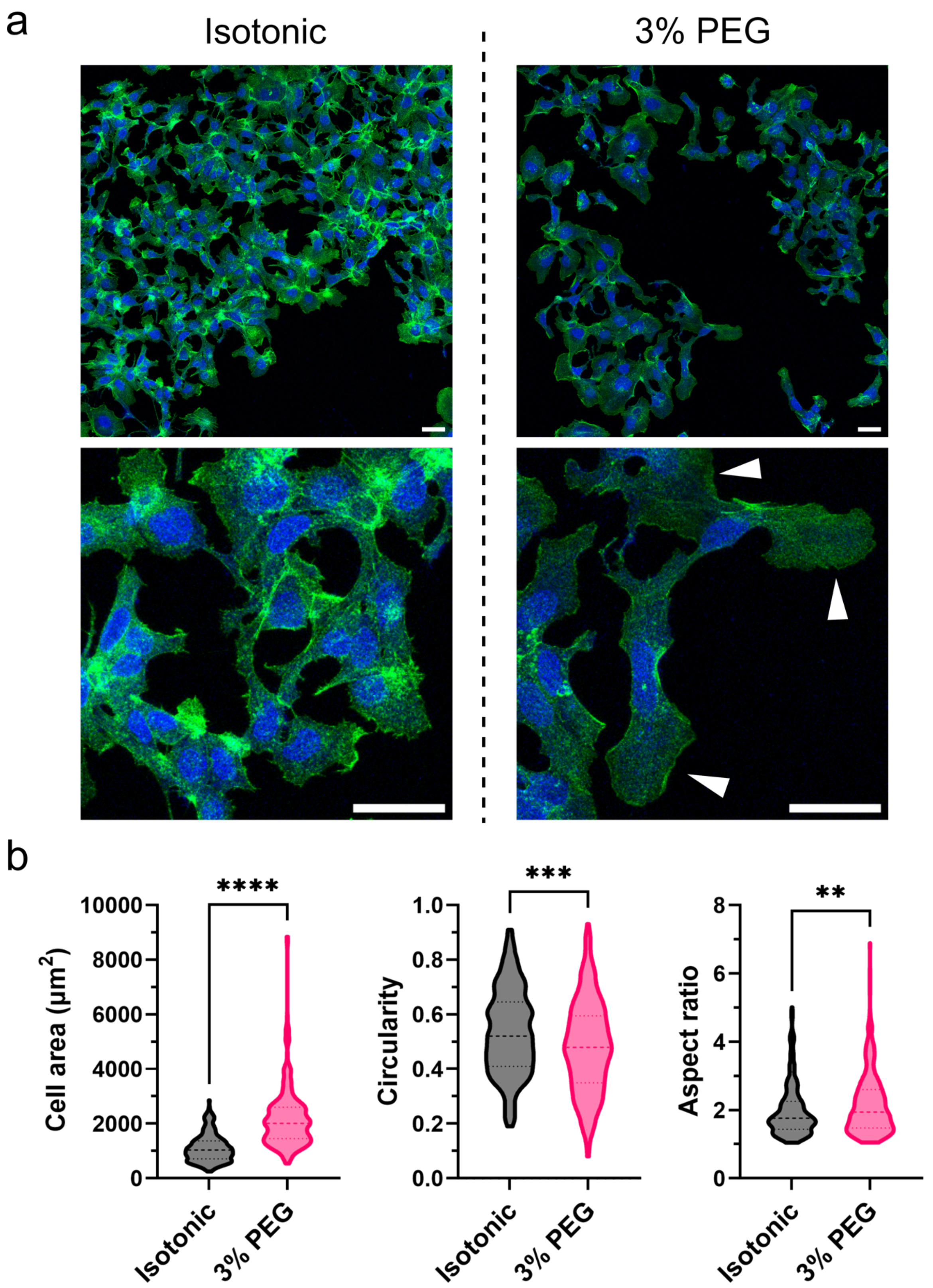

a
Isotonic
3% PEG
b
****
***
**
10000
8000
6000
4000
2000
0
Cell area (μm²)
1.0
0.8
0.6
0.4
0.2
0.0
Circularity
8
6
4
2
0
Aspect ratio
Isotonic
3% PEG
Isotonic
3% PEG
Isotonic
3% PEG

**Figure S7. Volumetric compression induced morphological changes in liver cancer cell line Hep3B.** (a) Fluorescence staining of the day-5 colonies under isotonic and 3% PEG conditions revealed: (b) a significant increase in cell spreading area and elongation, and reduced cell circularity. (n= 241 cells for isotonic condition, and 265 cells for 3% PEG condition, from N=2 biological replicates each condition). The arrowheads indicates the increased lamellipodia formation. Scale bars, 50 μm. A two-tailed student's t-test was used (**$p<0.01$; ***$p<0.001$; ****$p<0.0001$).

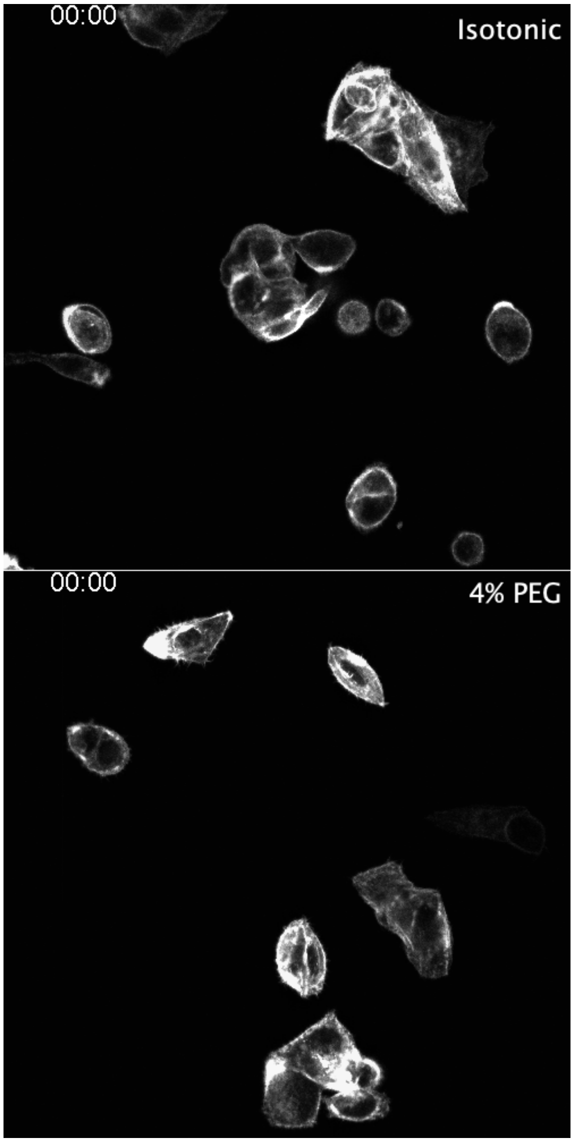


**Movie 1.** Time-lapse imaging (20 hours) of the morphology and actin dynamics of HepG2-LifeAct cells from Day 0 to Day 1 under 4% PEG compression compared to the cells cultured in the isotonic medium.

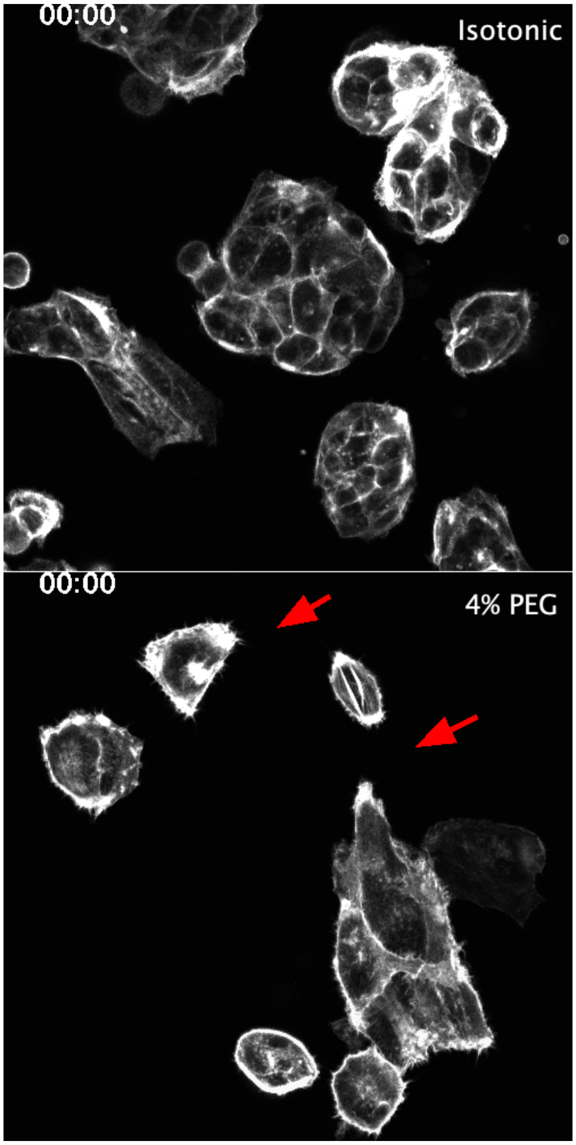


**Movie 2.** Time-lapse imaging (14 hours) of the morphology and actin dynamics of HepG2-LifeAct cells from Day 3 to Day 4 under 4% PEG compression compared to the cells cultured in the isotonic medium. The observed colonies are the same colonies monitored in Movie 1. Red arrows indicate the lamellipodia protrusions generated over the course of 14 hours.
